# A look beyond topography: transient phenomena of *Escherichia coli* cell division captured with high-speed in-line force mapping

**DOI:** 10.1101/2024.07.08.602450

**Authors:** Christian Ganser, Shigetaka Nishiguchi, Feng-Yueh Chan, Takayuki Uchihashi

## Abstract

Life on the nanoscale has been made accessible in recent decades by the development of techniques that are fast and non-invasive. High-speed atomic force microscopy (HS-AFM) is one such technique that has proven to shed light on elusive mechanisms involving single proteins. Extending HS-AFM to effortlessly incorporate mechanical property mapping while maintaining fast imaging speed allows us to look deeper than topography and reveal more details of the nanoscale mechanisms that govern life. Here, we present high-speed in-line force mapping (HS-iFM), which enables the recording of mechanical properties and topography maps with high spatiotemporal resolution. Employing this method, a detailed study of the dynamic nanoscale mechanical properties of living *Escherichia coli* bacteria reveals localized stiffening during division, intricate details of the division process, formation and diffusion of pores in the membrane, and the impact of depressurization of a cell. All of these phenomena were recorded with a frame time as low as 15 s and a spatial resolution of 5.5 nm/pixel in topography and 22 nm/pixel in force maps, allowing the capture of transient phenomena on bacterial surfaces in striking detail.

---

Atomic force microscopy (AFM) [1] employs physical interaction of a sharp tip at the end of a cantilever with a sample to generate images of a wide variety of materials beyond the optical resolution limit. While originally designed for surface science applications to overcome the limitation of scanning tunneling microscopy [2] to conductive materials, AFM was soon discovered to provide unique insights into biological samples [3–6]. To further the applications to biological samples – which are soft and dynamic in nature – conventional AFM was eventually found to be insufficient due to slow scan rates on the order of several minutes per frame and comparatively high invasiveness. To overcome these limitations, high-speed AFM (HS-AFM) was developed, enabling real-time imaging of fragile biological samples such as single proteins at high spatiotemporal resolutions [7]. Focusing solely on topography measurements to analyze structures and dynamics of proteins, unique insight into biological phenomena was gained [8–10]. Further applications to living cells [11,12] and bacteria [13] demonstrate that HS-AFM is highly flexible, recording not only single protein motions but also dynamics at the cell level – spanning several orders of magnitude in scale.

Recently, however, the potential of HS-AFM to go beyond topography measurements and extract mechanical properties has garnered attention [14]. In conventional AFM, methods to test mechanical properties were realized soon after its inception [15,16]. This mechanical testing with AFM opened up a whole new perspective on nanomechanical studies which delivers quantitative results by employing contact mechanics theories [17– 20]. Naturally, this included also biological samples [21–24] which are often too soft or fragile to be investigated otherwise. Nowadays, AFM based nanomechanical testing employing individual force curves to study elastic or plastic responses or full-fledged force mapping [25,26] comprised of hundreds or thousands of force curves are standard tools in the repertoire of nanomechanical investigations [27–33]. Such nanomechanical modes can be time consuming, increasing the time to record a single frame manifold compared to pure topographical modes, leading to the recent development of high-speed force curve measurements [34]. Recent reports mention force map techniques with frame times of 1.5 min [35] and 26 s [36], with a resolution of 128 px × 128 px and 64 px × 64 px, respectively. However, in both cases, topography resolution is dictated by force map resolution and increasing the number of pixels would inevitably increase the frame time. Promising alternatives to classical force mapping are so-called off-resonance techniques, which allow dynamic force measurements that can operate at high speeds and high resolutions [37,38]. Such methods are perfect for low force applications and when viscoelasticity is of little concern allowing for high frequency testing. In other cases, however, a lower force rate or larger forces are desirable to fully grasp the mechanical situation, which is better achieved by traditional force mapping.

Building on the previously introduced in-line force curve measurements [34] – which allow force curve acquisition without stopping the imaging process – we demonstrate the application of high-speed in-line force mapping (HS-iFM) to living *Escherichia coli* cells. *E. coli* are gram-negative bacteria and commonly found in the vertebrate gut microbiome. The wide use of *E. coli* as a model organism [39] makes them perfectly suitable for investigation with HS-iFM, because data such as AFM surface structure, mechanical properties [40–45] and general handling procedures [46] are readily available. In addition, *E. coli*’s doubling time is as low as 20 min at 37°C and still below 1 h at room temperature (∼25°C) [47–49], leading to a dynamic system that will be difficult to be investigated with conventional force mapping to capture transient phenomena. While the mechanical properties of *E. coli* cells are widely studied, their temporal evolution and impact on the life cycle remains elusive. During their life cycle, *E. coli* generate membrane vesicles, show dynamic membrane rearrangement, and – most obviously – constrict drastically in a highly localized fashion during division. Using HS-iFM combined with fluorescence microscopy, the division process of *E. coli* is visualized in striking detail, not only in terms of mechanical properties at the cell level but also with high-resolution topography images simultaneously capturing features on the scale of 10 nm. This unprecedented level of detail provides new insights into the dynamic nanoscale processes governing bacterial cell division and highlights the potential of HS-iFM in the study of living systems.

## High-speed in-line force mapping (HS-iFM)

HS-iFM is based on in-line force curve (iFC) measurements, which are designed to record a force curve during tapping mode without interrupting the scanning process, as shown in Figure 1a [34]. HS-iFM collects multiple iFCs arranged in a predefined grid, enabling to study spatiotemporal variations of mechanical properties. The principle of multiple inline force curves is explained in Figure 1b and 1c and it follows the same principle as for iFC measurements [34]. In short, the sample surface is scanned in tapping mode. The *x*- and *y*-positions are incremented until reaching one of the predefined force curve points, where a force curve (Figure 1d) is recorded, e.g. (*x*_*0*_,*y*_*0*_) indicated in Figure 1e. At this point, the position is held constant while force curve acquisition is completed and scanning resumes until the next point (*x*_*1*_,*y*_*0*_). This procedure continues until the last point (*x*_*n*_,*y*_*n*_) is reached. This process is substantially different from conventional AFM force mapping, where topography is reconstructed from the force curves and thereby inextricably linking the resolution of topography maps with force maps. In HS-iFM, however, topography is recorded in tapping mode within the same frame as the force map by rapidly switching between force and topography measurements, allowing to choose topography pixel resolution independently from the force map. A detailed explanation of HS-iFM and the signals involved is presented in Supplementary Note 1 and Supplementary Figure S1, an analysis of the frame time and variables influencing it is given in Supplementary Figure S2, and removal of stripe artifacts caused be force mapping is shown in Supplementary Figure S3.

**Figure 1:**
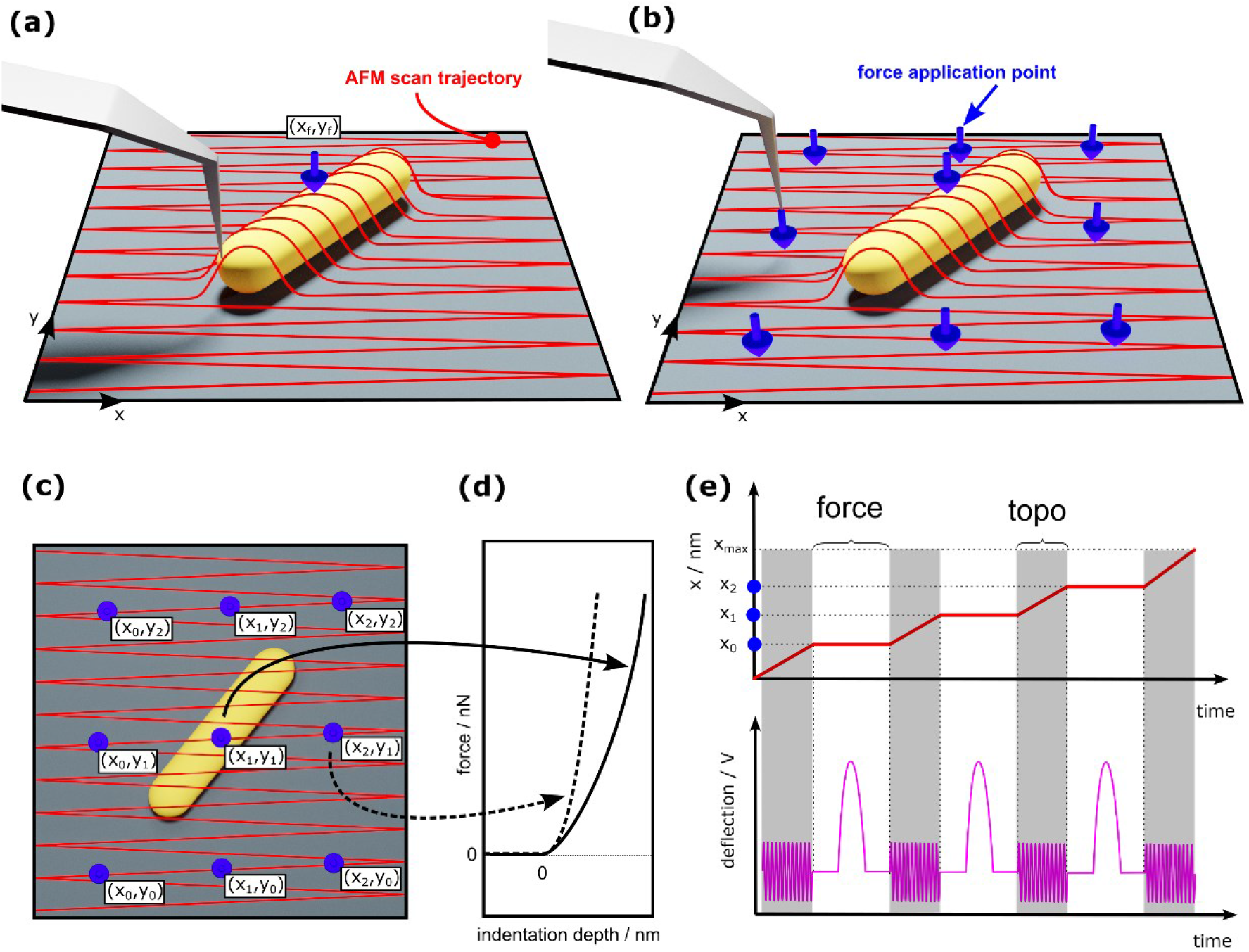
Schematic representation of HS-iFM. (a) In-line force curve measurements are the basis of HS-iFM but record only a single force curve per frame, indicated with the blue arrow and the coordinate (x_f_,y_f_). (b) In HS-iFM, multiple force curves arranged in a predefined grid are recorded per frame. The blue arrows indicate an exemplary 3 × 3 force map. (c) Top view of (b) with all the force application points marked by coordinate pairs. (d) Force curves extracted from two points in (c) from areas with different mechanical properties. (e) A scheme of the x-axis motion as well as cantilever deflection over time. Every time a force coordinate is reached (e.g. x_0_, x_1_, and x_2_), scanning is paused and a force curve is recorded. Force curves are captured in the “force” regions, while in the regions marked “topo” topography is recorded and scanning is resumed.

## Dynamics of mechanical properties during *E. coli* division

A demonstration of continuous recording of HS-iFM on living *E. coli* is presented in Figure 2. The division process of a cell into two daughter cells was successfully observed simultaneously in topography and mechanical properties (Figure 2a) as well as fluorescence imaging (Figure 2b). Filtered topography maps, generated using high-pass filtering of original topography maps, revealed detailed structures of the cell’s surface. The initial snapshot (I) shows that the initial surface of the bacterium is smooth without the visible division site, and the elastic modulus map exhibits no remarkable features except for a slightly stiffer bottom part compared to the top part. Furthermore, in the topography holes are visible that close up and reform again (marked by green arrows), which are also discernible in the modulus map as slightly softer parts on the surface. This dynamic hole formation could be attributed to the formation of outer membrane vesicles, which are reported to be more frequent during cell division and around septa [50,51]. In image (II) of Figure 2a, the same two former holes have become bulges and are joined by another bulge between them. Interestingly, all three bulges are pinned to the division site and remain softer than the surrounding area. In the modulus map, the division site has become visible as a straight line separating the softer top part from the stiffer bottom part. The simultaneously recorded fluorescence image in Figure 2b shows a decreased intensity in the center of the cell. This phenomenon is caused by the incorporation of new cell wall material around the division site. The cell wall material was chemically labeled by Alexa-488 WGA before the measurement and no fluorescent dyes were present in the medium, thus, the new cell wall components appear dark. From here on, the division site becomes increasingly stiffer, while the upper and lower part of the dividing cell do not change significantly (Figure 2c). At (III) in Figure 2a, the division site is clearly visible in both modulus and topography, while the bulges are seen only in topography because they have shrunk and have less impact to the local modulus. The division continues to stiffen until (IV), where significant contraction of the cell is already evident. From this point on, the division site softens again, while both upper and lower cells become stiffer until a plateau of about 0.75 MPa is reached. At position (V), the division site is no longer recognizable as stiffer in the modulus map and from the topography it is plausible that at this point, the bacterium has finished dividing and was separated into its two daughter cells. Continued growth of the daughter cells is observed until (VI) where it became unambiguously clear that they had separated. Interestingly, after the separation, the bottom cell becomes significantly stiffer, while the top cell does not. The division process in its entirety is shown in Supplementary Video 1.

**Figure 2:**
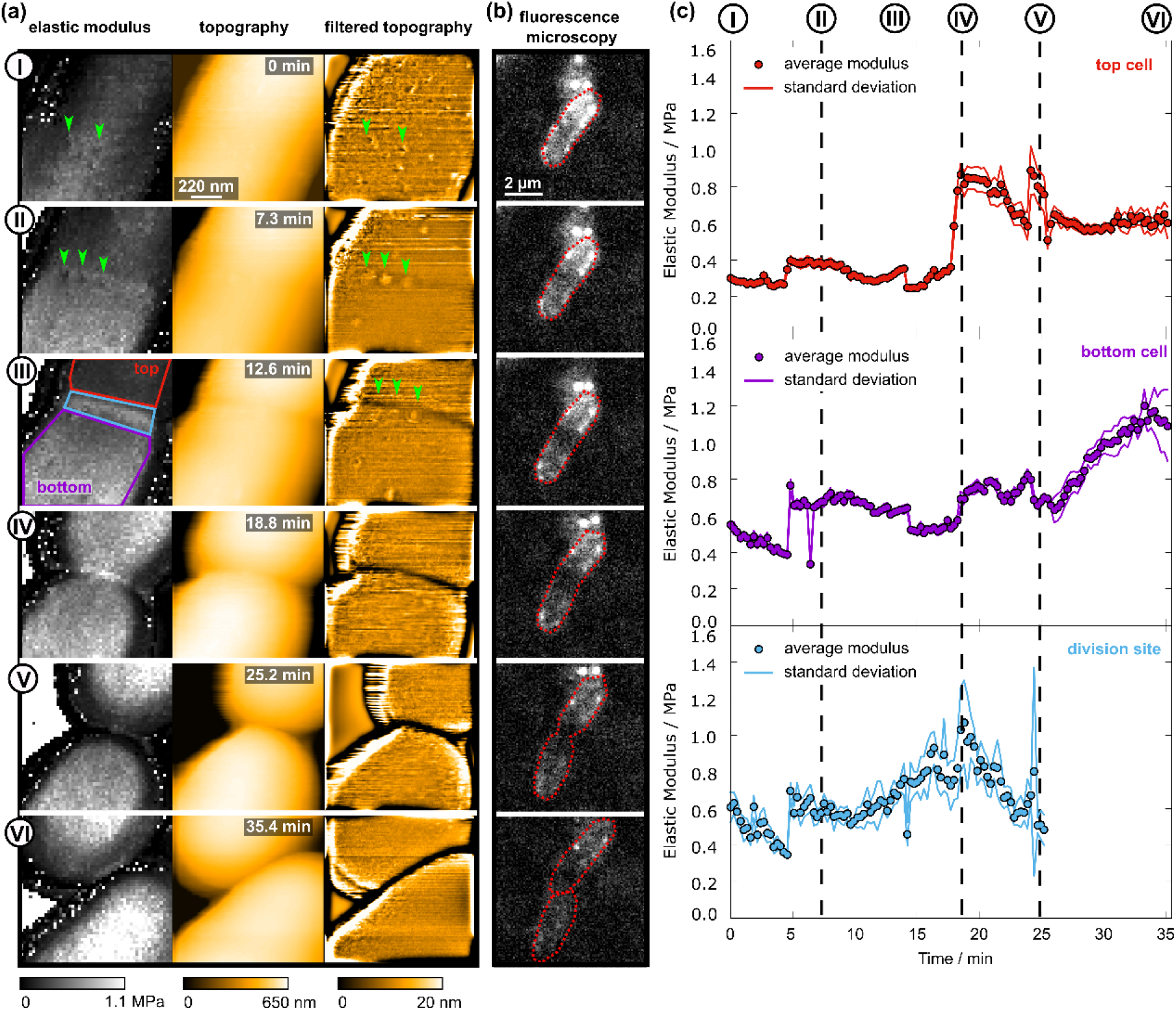
Elastic modulus of E. coli during cell division. (a) Snapshots of the elastic modulus and HS-AFM topography during several stages of cell division. The rightmost column shows the bacteria surface topography after high-pass filtering to remove the surface curvature. (b) Fluorescence microscopy images recorded simultaneously with the HS-AFM images. (c) Temporal evolution of the elastic modulus of the top cell, the bottom cell, and the division site, respectively. The points are the average values of the respective areas and the solid lines show the standard deviations, determined according to Supplementary Figure S4. The imaged area is 1.1 µm × 1.1 µm, the time to record a single frame was 16.1 s. Topography has a resolution of 150 × 150 pixel, while modulus maps have a resolution of 50 × 50 pixel.

## Mechanical considerations during cell division

HS-iFM measurements revealed an increasing stiffness of the division site which could be a consequence of the division mechanisms of *E. coli*. However, before drawing any conclusions based on the measured elastic modulus values, it is necessary to exclude topographical artefacts that could impact the contact mechanics analysis. Due to the highly concave surface at the division site, the effective contact area of the tip-sample interface is increased, potentially leading to an overestimation of the elastic modulus. To confirm whether the topography has any effect on the measured elastic modulus, Hertzian contact mechanics was employed.

First, the radius of the division site was determined longitudinally and perpendicular to the cell’s long axis. The longitudinal radius was measured along 9 parallel lines centered around the long axis (Supplementary Figure S4). From each line, the height profile was extracted and was fitted with a parabola at the minimum point. From these 9 fits, the lowest radius was selected to ensure that the highest possible influence would be determined. Subsequently, a perpendicular line profile at the bottom of the division site was fitted with a circle to obtain the perpendicular radius. This process was repeated for all frames recorded on the cell from 0 min to ∼25 min (95 frames). The results are presented in Figure 3a and the relevant radii are illustrated in Figure 3b. Note that the longitudinal radius, in particular, will be influenced by tip-sample dilation and will appear lower than it actually is. To correct this, the assumed tip radius of 5 nm was added to measured longitudinal radius (Supplementary Figure S5). It is evident that both longitudinal and perpendicular radius decrease with progressing division.

**Figure 3:**
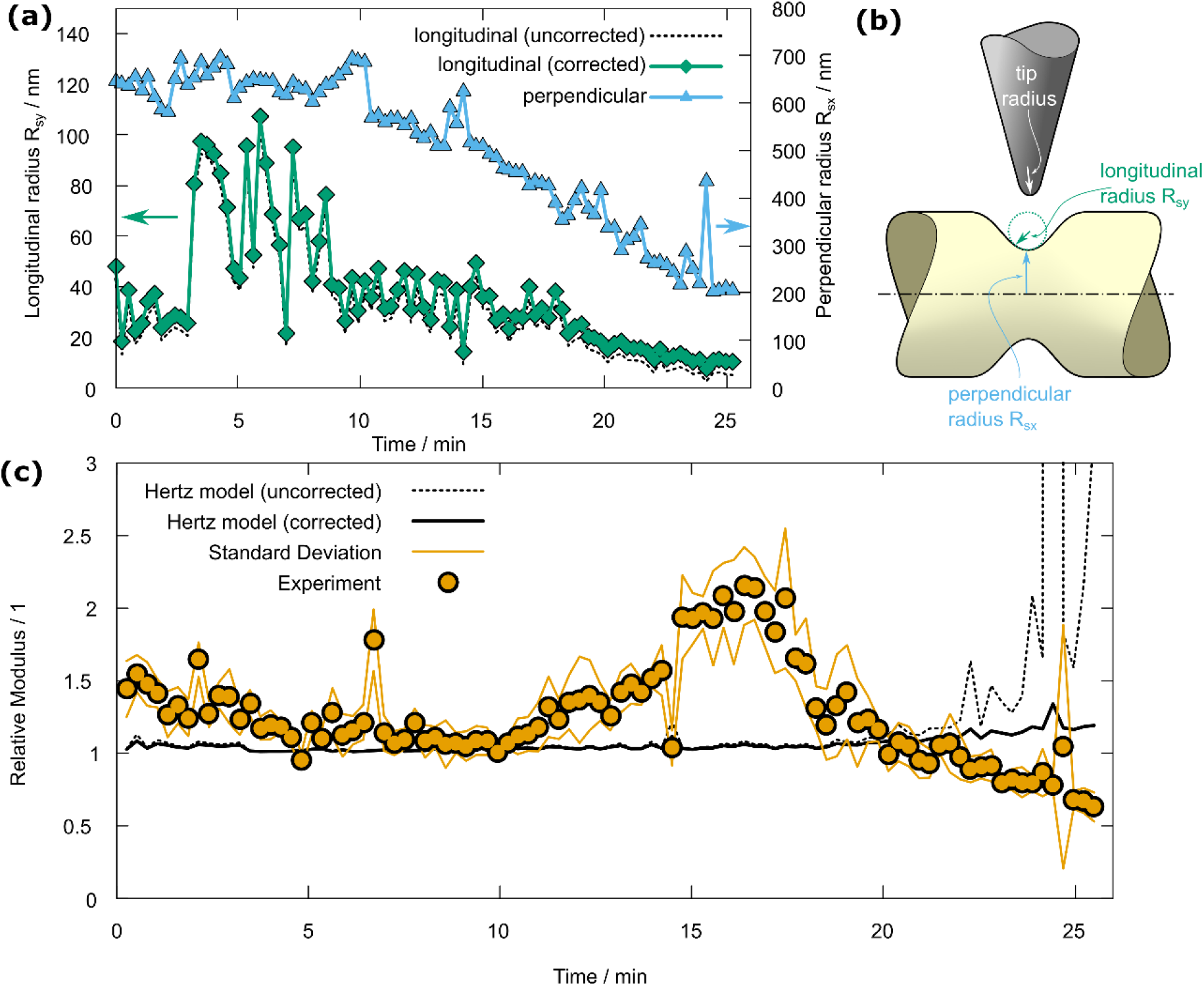
Contact mechanical considerations of tip-sample interaction at the division site. (a) Development of the radius on the bottom of the division site in the cell’s longitudinal (green line), R_sy_, and perpendicular direction (blue line), R_sx_. The black dashed line indicates the longitudinal radius without correction for tip-sample dilation. Until about 10 min, the division site was not formed and R_sy_ is likely underestimated and dominated by local surface roughness causing the fluctuations seen in the plot. (b) Illustration of longitudinal, perpendicular, and tip radius. (c) Elastic modulus of the division site relative to the area outside of the division site (orange) compared with the expected increase due to topography (black lines). The solid line was calculated using the Hertz model with the corrected radii from (a) and assuming a tip radius of 5 nm. The dashed line is the result when using the uncorrected longitudinal radius from (a).

Next, the expected relative apparent increase of elastic modulus due to the topography was calculated using Hertzian contact mechanics with an elliptical contact area. While the tip is assumed to have rotational symmetry, the same cannot be assumed for the bacteria. Before the division starts and on areas far from the division site the cells’ radius is around 600 nm which is significantly larger than the tip radius (assumed to be 5 nm) and the longitudinal radius is effectively infinite. However, during division the pronounced localized constriction of the membrane not only causes the cell radius to decrease but also creates a deep valley around the circumference with a radius at the bottom that approaches the tip radius as time progresses (Supplementary Figure S6 and S7). Generally, such a case will lead to an elliptical contact area – in contrast to the circular contact area in the classical Hertzian theory. This necessitates the use of a more generalized Hertzian contact theory [52] which allows for a correction factor *f(e)* as a function of the contact area’s eccentricity *e*. The eccentricity will depend on the tip radius and the sample curvature radius in *x* and *y* direction, *R*_*sx*_ and *R*_*sy*._. The full calculation is described in detail in Supplementary Note 2.Using the cell’s radii from Figure 3a so that *R*_*sx*_ is the perpendicular radius and *R*_*sy*_ is the longitudinal division site radius, *f(e)* can be plotted for each frame and is presented as the black solid line in Figure 3b. The factor *f(e)* is compared to the modulus of the division site relative to the average modulus of the cell outside of the division site, presented as solid orange circles line in Figure 3b. It is quite obvious that the topography change has only minimal effect and cannot account for an increase in modulus by a factor of 2. Even if the assumed tip radius is increased to 100 nm, such a stark rise in elastic modulus is not likely to be caused by topographical artefacts (Supplementary Figure S8). The dashed line in Figure 3b is also *f(e)* but calculated with the dilated *R*_*sy*_ from Figure 3a. It is clear that even in this case, the curvature will not significantly influence the apparent elastic modulus. Only when the division site has almost disappeared, can a drastic increase of the uncorrected *f(e)* be observed.

In conclusion, it is fair to assume that the measured increase of the elastic modulus on the division site is, indeed, real and not caused by the significant topographical changes. Similar effects as described above, such as stiffening of the division site not being influenced by topography, were commonly observed on dividing bacteria (Supplementary Figures S9 – S11, Supplementary Videos 2 – 4).

Another mechanical observation concerns the dynamic changes in elastic modulus. In Figure 2a (II), the bottom cell appears stiffer than the top cell. Supplementary Video 4 shows a similar case with the soft part detaching from the substrate after division, suggesting partial cell attachment. This is important because cell attachment is not fully controllable, potentially leading to misinterpretations in conventional force mapping and individual force curve measurements. HS-iFM combined with optical microscopy can identify such issues through time-resolved observations. It is advised to consider the entire cell and be cautious with gradients or abrupt changes in elastic modulus. In Figure 2, the top cell’s initial modulus is underestimated while the bottom cell is mostly unaffected. The division area’s modulus might be underestimated, but the relative modulus of the division site in Figure 3b shows a clear increase in stiffness. Without HS-iFM’s spatiotemporal resolution and the simultaneous optical microscopy imaging, these effects might be not be detectable at all. The reliability of modulus values presented is confirmed when compared to widely available results from AFM-based investigations. Our measurements result in elastic modulus values ranging from about 0.8 MPa to 2.5 MPa, excluding unbound areas and division sites, which compares well to reported values of 3.5 MPa ± 1.6 MPa [45], 0.5 MPa to 1.7 MPa [42], 1 MPa to 10 MPa [41], and 6 MPa ± 3 MPa [44]. All but the latter values were obtained from *E. coli* in liquid environment. A notable exception is an elastic modulus of over 200 MPa [43], obtained from individual force curve measurements with stiff cantilevers on dried *E. coli* in air.

From the contact mechanical considerations, geometrical effects play no significant role in the stiffening of the division site. However, there are two mechanisms crucial to bacterial cell division that have the potential to cause localized stiffening of the division site. First, the initiation of cell division is governed by the formation of the so-called Z-ring [53], which initiates mechanisms that cause the membrane to curve inward and constrict [54,55]. This inward pulling of the cell wall causes a significant amount of stress, especially at the bottom of the division site where pressure builds up [55]. This stress will cause the membrane to stiffen locally and lead to an increased elastic modulus as seen in Figure 2 and Supplementary Figures S9 – S11. The mechanical properties of the membrane do not change, but the increased stress causes a stiffer response of the membrane to the indenting tip [56,57]. The second factor that contributes to the stiffening of the division site is the gradual formation of the septum. The abovementioned constriction leads to an invagination of the inner membrane and the formation of a peptidoglycan wedge between the outer and inner membranes [58]. This wedge continues to grow until it forms the septum. It stands to reason that such a wedge will have increased structural stability and thus increase the measured elastic modulus. Using HS-iFM, we directly demonstrated that dynamic stiffness increase of the division site is a characteristic feature in *E. coli* cell division.

## Nanoscale features during cell division

In addition to mechanical property measurements, HS-iFM also facilitates high-resolution topography imaging, which is presented in Figure 4a, as well as Supplementary Figure S11. Close inspection of such high-pass filtered high-resolution topography images, which are recorded simultaneously during force mapping revealed pores on the surface of *E. coli* cells (Supplementary Video 5). These pores dynamically close up (Figure 4b) and appear to diffuse (Figure 4c). A possible explanation of these pores could be outer-membrane protein complexes located on the bacteria surface [6,59]. The diameter of the pores was measured to be 34.7 nm ± 11.8 nm (average and standard deviation of 8 pores, Supplementary Figure 12). This is noticeably larger than what was reported for protein complexes, which are approximately 8 nm in diameter and arranged in a fairly regular pattern [6]. Because the pores found here are mostly isolated and are dynamic in nature, they could be attributed to dynamic rearrangement of the outer membrane and related to membrane vesicles.

**Figure 4:**
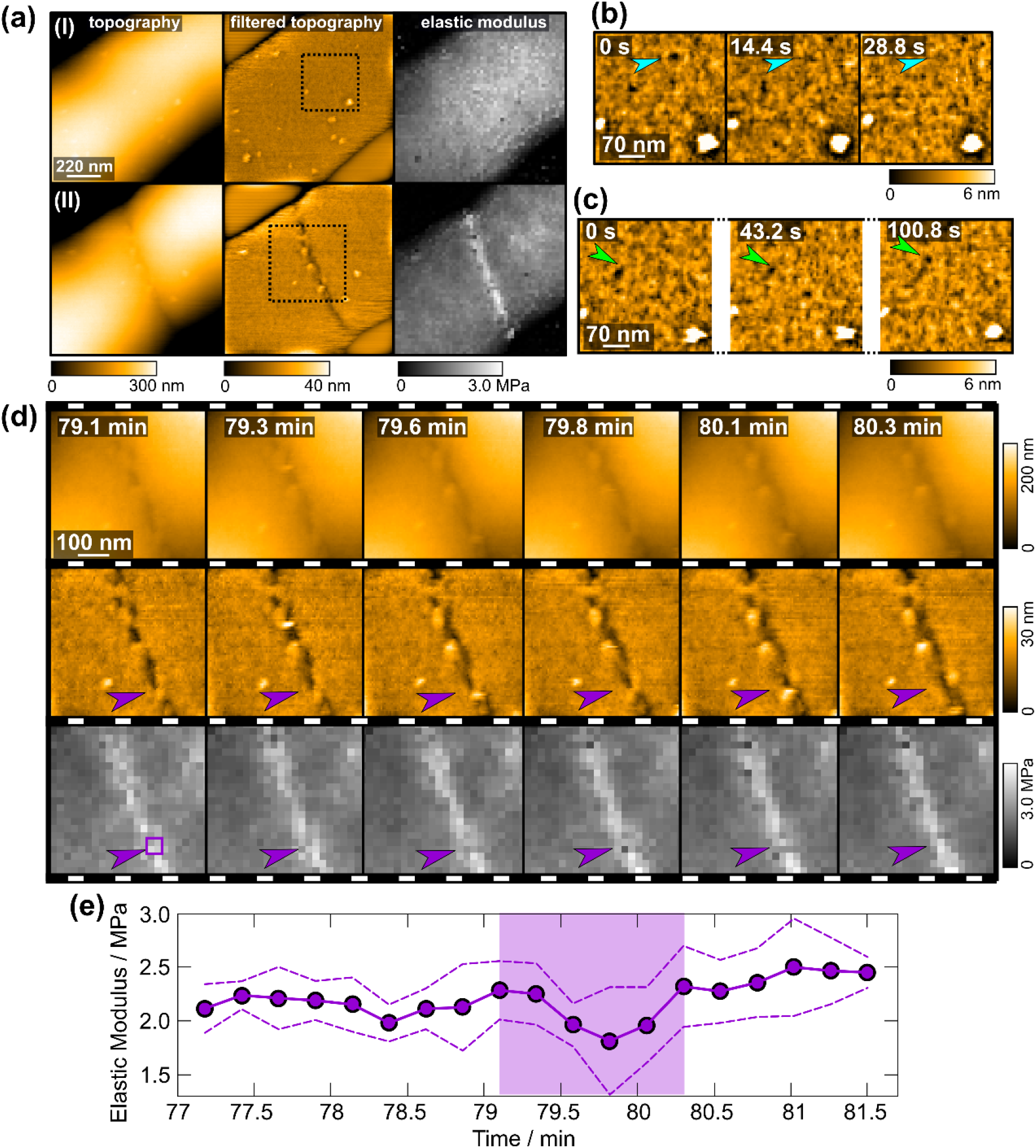
High resolution imaging of E. coli cell division as seen in Supplementary Video 4. (a)Topography, filtered topography, and elastic modulus map at the beginning (I) and in the middle (II) of cell division. (b) Zoomed sequence of the area marked in (a)-(I) showing a pore (indicated by cyan arrows) closing. (c) Pore diffusion (indicated by green arrows) observed in the area marked in (a)-(I). (d) Details of cell division observed in the area marked in (a)-(II) as topography (top), filtered topography (middle), and elastic modulus (bottom). The purple arrow points to a bridge that is present during division and brakes as the cells continue to separate. The elastic modulus softens locally when the bridge brakes. (e) Elastic modulus average plot from the area indicated by the purple 4-pixel quadrilateral in (d). The purple tinted area corresponds to the images shown in (d), where a small dip in modulus is visible. The full imaged area as shown in (a) is 1.1 µm × 1.1 µm, the time to record a single frame was 14.4 s. Topography has a resolution of 200 × 200 pixel, while modulus maps have a resolution of 50 × 50 pixel.

Further inspection of the division site at high resolution also revealed striking details of the division process. During division, the membrane appears to form bridges between both cells that are stretched and subsequently ruptured (Figure 4d). These bridges become softer while rupturing (Figure 4e) and after the bridge has broken down, the original stiffness is recovered. This process is reminiscent of the formation of outer membrane vesicles, as the bridge also forms a small bubble that disappears promptly. HS-iFM with high spatiotemporal resolution has proven not only to offer mechanical insight into bacterial cell division, but also link it with nanometer scale dynamics of the division site and record membrane processes in striking detail.

## Cell depressurization

In a serendipitous moment, a sudden loss of cell rigidity was observed during continuous HS-iFM measurements as is presented in Figure 5, Supplementary Video 6, and Supplementary Figure S13. In Figure 5a, topography, elastic modulus maps and optical microscopy images are shown for different times during HS-iFM observation and a zoom of the area marked with the blue arrow is presented in Figure 5b. The evolution of the elastic modulus is plotted in Figure 5c, with the positions of the images marked as (I) – (IV). An *E. coli* cell (I) starts to grow and develops the stiffening of the division site (II). The division progresses (III) and a clearly soft substance coats the division site obscuring it completely. At the 52 min mark the elastic modulus of the dividing cells drops suddenly and drastically by a factor of 4 (from about 0.8 MPa to 0.2 MPa) and remains low until the end of the observation (IV). During most of the time, a separate cell, defined as reference cell, was partly visible in the bottom right corner (marked with a white arrow). It is noteworthy that the elastic modulus of the reference cell does not drop and remains largely unchanged, as can be seen in Figure 5c. The fact that the second cell does not soften excludes a measurement artefact, e.g. a sudden change of the tip due to contamination. Furthermore, the loss of stiffness is concurrent with the formation of a blob on the cell surface (marked with a blue arrow in Figure 5a). The blob’s elastic modulus is close to 0 Pa, suggesting that it is too soft for meaningful determination of mechanical properties. This blob originates from what appears to be a membrane vesicle, which is present from the very beginning of the observation (see also Supplementary Figure S13). Another blob is visible at the division site in Figure 5a (III) and (IV), which forms at about 37 min (see Supplementary Video 6), more than 15 min before the loss of stiffness and seems not to be directly related to the cell bursting. The vesicle might be a weak point in the cell that ruptures during imaging, releasing the soft protoplasm upon bursting. Puncturing of the cell causes the turgor pressure to be released, equilibrating the inside of the cell with the environmental pressure which leads to a reduction in membrane tension and subsequently to a reduced elastic modulus. Therefore, the elastic modulus after depressurization can be attributed purely to the cell wall and its structural configuration, e.g. the shape of the cell. Despite only the left side being punctured, both sides depressurize simultaneously, indicating that septation was incomplete and the cells’ interiors were still connected. Such observations have the potential to provide unique insight into the cell division cycle by identifying when septation finishes and the daughter cells’ interiors are no longer connected.

**Figure 5:**
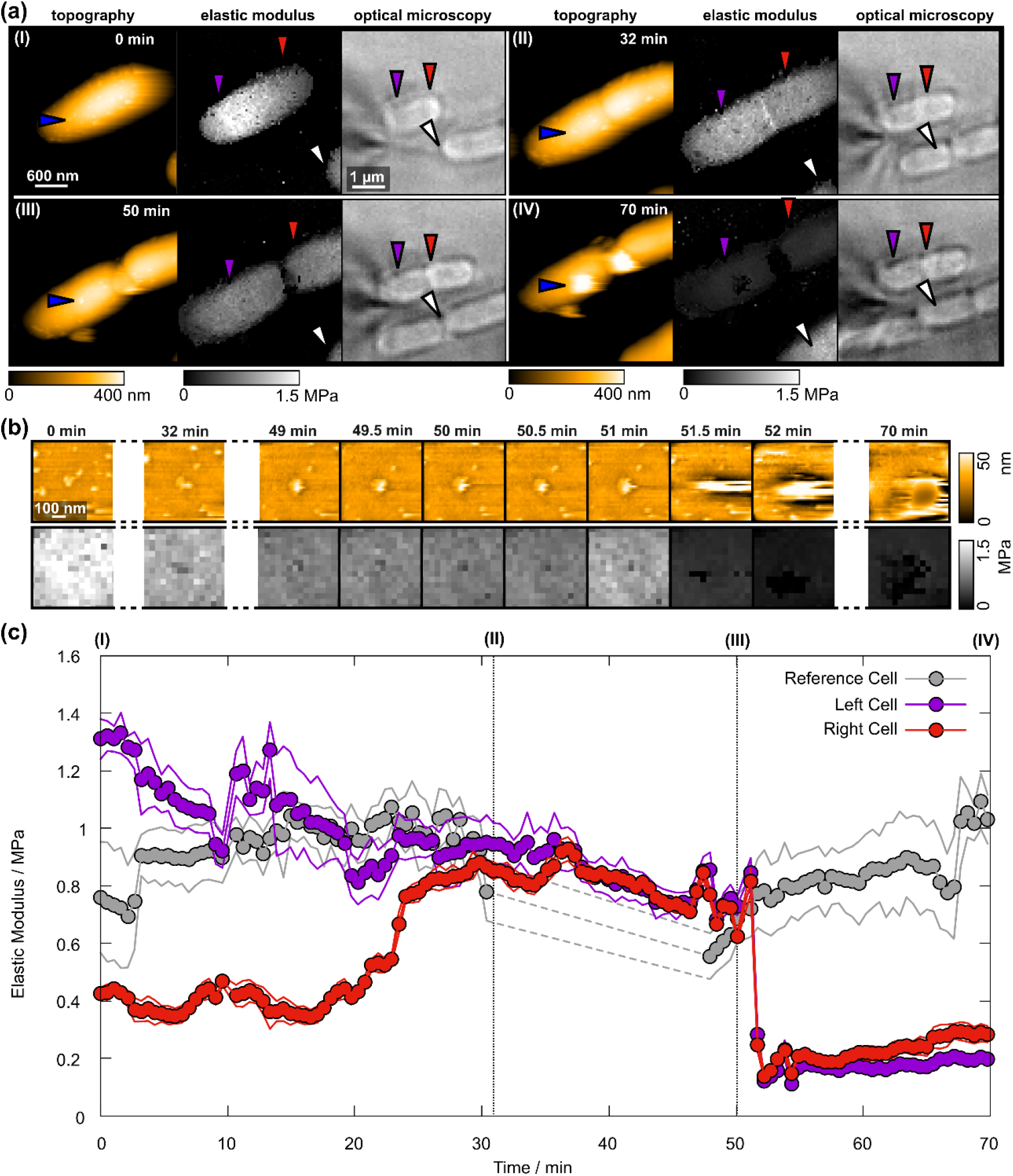
Bursting E. coli cell during HS-iFM imaging. (a) Topography, elastic modulus, and optical microscopy images of a living E. coli cell which grows and starts dividing (I)-(III) before bursting (IV). Purple and red arrows mark the left and right cells respectively, while the blue arrow marks the spot where the cell bursts. The white arrow marks a nearby cell which retains its stiffness for reference. (b) Zoom-ins of the area around the bursting site of topography and elastic modulus. (c) Time evolution of the elastic modulus of the left (purple) and right (red) sided of the cell compared with a reference cell (gray). The lines mark the standard deviations. Between 30 min and 47 min the reference cell was outside of the observation area. The elastic modulus of the left and right cell was determined the same as in Figure 2 (described in Supplementary Figure S4). The elastic modulus of the reference cell was averaged over the visible portion of each frame.

## Conclusions

The application of HS-iFM to observe the life cycle of living bacteria reveals striking details at high resolutions in both topography and elastic modulus. Essentially, by using HS-iFM, it is not necessary to sacrifice high-resolution imaging capabilities for a comprehensive mechanical characterization at fast imaging speeds.

In this study, we demonstrated not only the measurement of local stiffening of the division site, but also – at the same time – dynamic hole formation, outer membrane vesicles, and detailed bridge formation and rupturing during division. The focus was mainly on larger area imaging to visualize the division in its entirety, however, there is great potential for imaging on a smaller scale as is evident by the observation of pores on the order of 30 nm. This may pave the way to actively image membrane proteins in living cells – not only in topography but gaining a thorough understanding of the mechanics involved in life on the nanoscale.

A bursting cell in the middle of the division process showed that puncturing the membrane leads to a loss of stiffness across the cell. This clearly demonstrates that a large part of the cell’s stiffness is caused by internal turgor pressure. However, some stiffness remains, which would largely be caused by the membrane’s mechanical properties and its structure. While a serendipitous observation, it alludes to greater capabilities – e.g. by locally increasing the force during HS-iFM to target weak points in the membrane and causing cells to burst by choice. This would open the possibility to directly study the effect of internal pressure on cell mechanics or to analyze the division cycle by investigating at which point septation has concluded and daughter cells have no internal connection remaining.

## Materials and Methods

### High-speed atomic force microscopy

The HS-AFM used for this study is laboratory-built and of the tip-scanning type, combined with an optical microscope, capable of fluorescent microscopy imaging. [60] Two scanners were available: a high-resolution scanner with a scanning range of 1.1 µm × 1.1 µm and a wide-area scanner with a scanning range of 8.3 µm × 8.3 µm. Scans with dimensions of 1.1 µm × 1.1 µm were recorded with the first scanner, while 3 µm × 3µm scans were recorded with the second scanner. The cantilevers in use were Olympus AC10 with a beak-like end on which a carbon tip was grown by electron beam deposition and sharpened by plasma etching. The tip radius is typically around 4 nm to 5 nm after plasma etching [61]. The manufacturer’s nominal spring constant for AC10 cantilevers is stated as 0.1 Nm^-1^ with a variation from 0.02 Nm^-1^ to 0.2 Nm^-1^. In order to precisely calculate the forces, however, the spring constant of each cantilever needs to be calibrated before each measurement. To do so, the thermal noise method was employed [62]. The cantilevers’ sensitivities – necessary to convert the deflection signal from a voltage to a physical distance – were determined by recording an HS-iFM map containing 2500 force curves on a glass surface before and after a measurement. For each force curve, the slope after contacting the surface was extracted by a linear fit. The histogram of the slopes was then fitted with a Gaussian distribution. The sensitivity was selected as the location of the Gaussian’s maximum value. The cantilevers’ average sensitivity was 20.7 mV/nm ± 0.9 mV/nm. The spring constants of the cantilevers were then found to be 0.11 N/m ± 0.054 N/m. Resonance frequency and Q-factor in the observation buffer were 475 kHz ± 29 kHz and 1.50 ± 0.12, respectively. All values above are given as mean value ± standard deviation from 11 cantilevers.

Topography scanning was exclusively performed in tapping mode, with the cantilever oscillating close to its resonance frequency. The free oscillation amplitude was set to be around 5 nm and the set-point during scanning was approximately 80% of the free amplitude.

For HS-iFM, the force map size was either 50 × 50 or 75 × 75 pixel, which provided sufficient resolution for spatial mapping of bacteria. Force curves were recorded with 1000 datapoints and a total time of 2 ms per curve.

The elastic modulus values were determined by fitting the force curves with the Hertzian contact model [17,63], which was selected because no adhesion forces could be detected in the force curves (see Supplementary Figure S14). The Hertzian model was designed to estimate the contact of spherical bodies without adhesion by approximating them with parabolas and is, thus, also valid for small deformation of spherical bodies [52,63,64]. However, for parabolic bodies, e.g. a parabolic tip indenting an elastic half space [18], the solution is exact and not limited for small deformations – as long as the deformations remain in the linear elastic regime. Due to the large number of force curves recorded, manual fitting of the data is not feasible and an automated approach is used (see Supplementary Figure S15).

### Sample preparation

Slide glass (Cat# C024321; Matsunami Glass) was covered by hydrophobic Teflon tape (Cat# ASF-110FR; Chukoh Chemical Industries) fixed with epoxy glue (Cat# AR-R30; Nichiban). A hole with a diameter of 5 mm was punched out in the center of the tape before gluing, to create a confined area for sample observation. Poly-L-lysine (70 kDa – 130 kDa, 1 mg/mL) was then added to the central glass surface and incubated for 5 min before washing it two times with 200 µL of milli-Q water. *Escherichia coli* (NBRC3972/ATCC8739) samples were purchased from the Biological Resource Center, National Institute of Technology and Evaluation (NBRC). The bacteria were cultured in lysogeny broth (LB) medium (Cat# 24308-08; Kanto Chemical) one day before the HS-iFM measurements. On the day of the measurements, the bacteria were stained by WGA Alexa Fluor^®^ 488 conjugate (Cat# W11261, Invitrogen) for 30 min. The stained bacteria suspended in LB medium were diluted by PBS to contain 10%-vol of the bacteria suspension. To immobilize the cells for the measurements, 2 µL of the diluted suspension was added to a 20 µL PBS droplet placed on the glass surface, diluting sample and LB medium to 1%-vol. The suspension was incubated for 10 min on PLL-glass to ensure that bacteria cells were securely immobilized. Higher concentrations of the sample lead to particles which are contained in LB medium to compete with the bacteria for adsorption, resulting in weak binding of the cells to the glass surface. After incubation, the glass slide was washed three times by 200 µL 10%-vol LB medium in PBS (observation buffer). Finally, the HS-iFM observations were performed in 200 µL observation buffer.

## Acknowledgements

This work was supported by JSPS KAKENHI Grant Number JP23H04874 in a Grant-in-Aid for Transformative Research Areas “Materials Science of Meso-Hierarchy” to C.G and 24K01309, 22K18943 to T. U and by JST, CREST Grant Number JPMJCR21L2, Japan to T. U.

We thank Kohei Fujihara (Nagoya University) for help with testing the implementation of HS-iFM.

## Supplementary Information for

### Supplementary Figures

**Figure S1:**
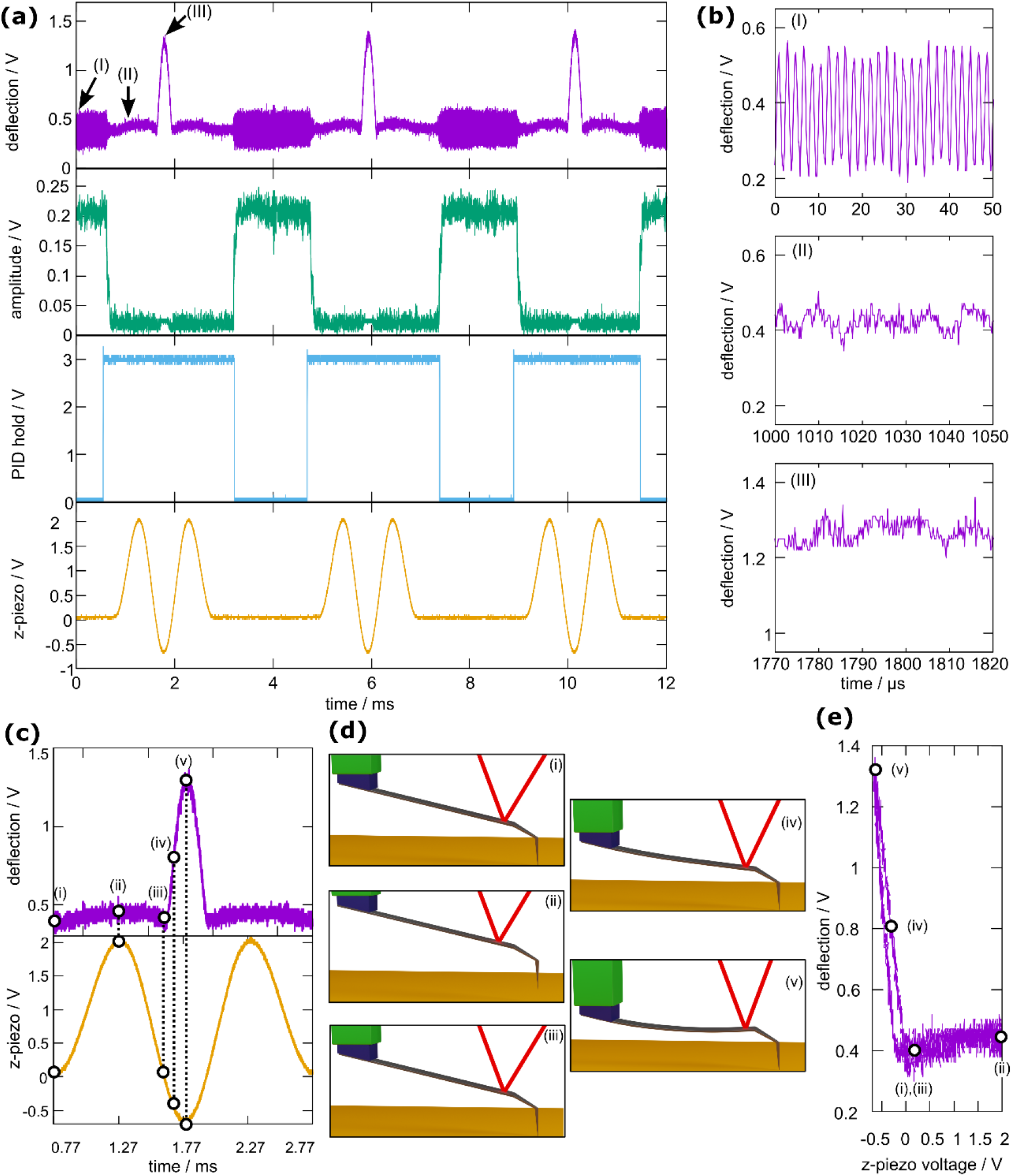
Signals during HS-iFM measurement. (a) Signal traces during force mapping showing three force pulses. The deflection signal shows the cantilevers raw deflection. The amplitude signal represents the cantilever’s oscillation amplitude at the excitation frequency. The PID hold signal switches the PID control on and off. Finally, the z-piezo signal is used to move the cantilever to apply forces to the sample.(b) Zoom-ins of the deflection signal at the locations marked in (a). (c) Enlarged deflection and z-piezo signals. (d) Cantilever states at the positions marked in (c). (e) Raw force curve gained by plotting the z-piezo voltage vs. the deflection shown in (c).

**Figure S2:**
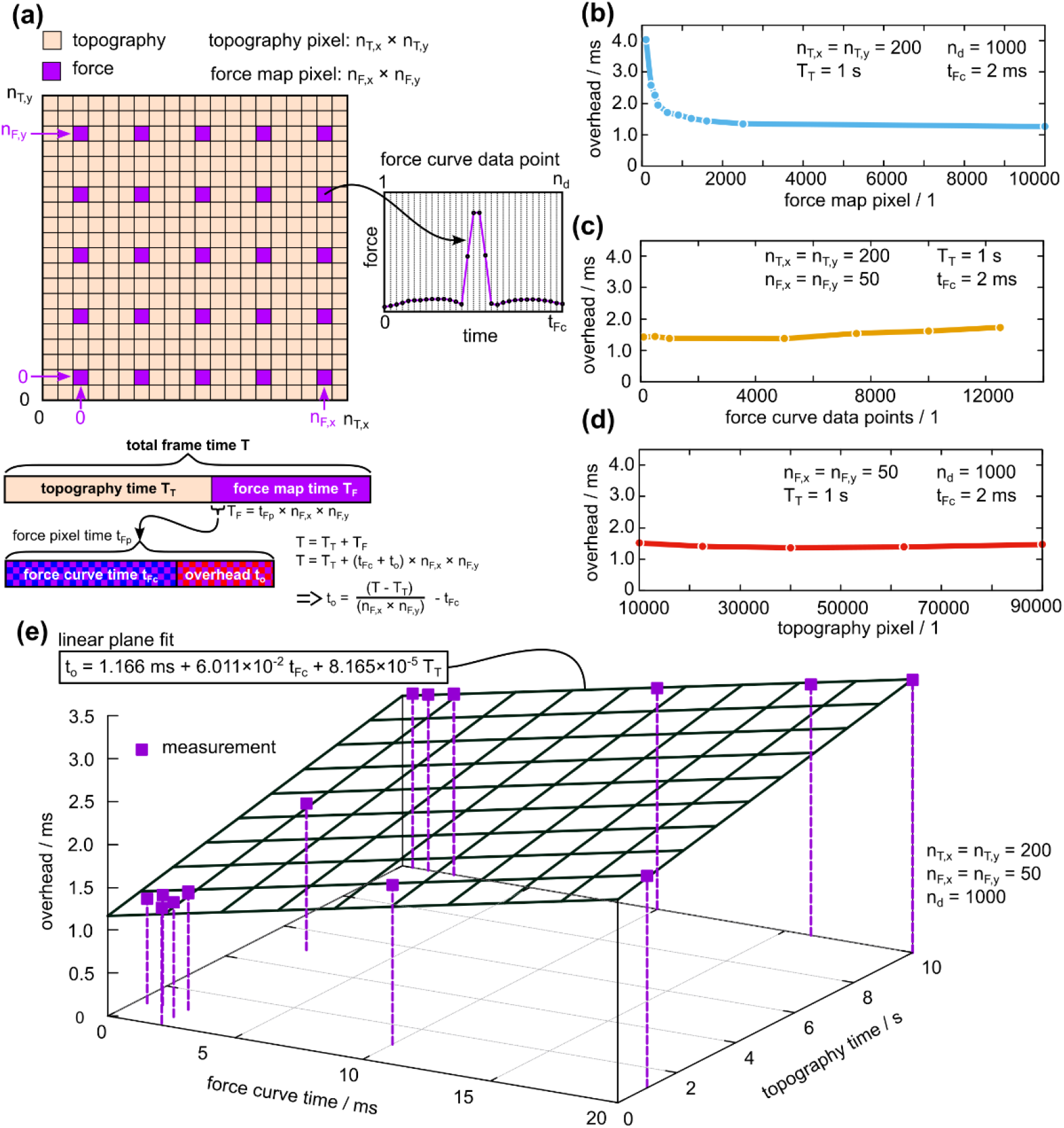
Frame time analysis during HS-iFM. (a) Graphical explanation of variables that can be freely set for and HS-iFM measurement (*n*_*T,x*_, *n*_*T,y*_, *n*_*F,x*_, *n*_*F,y*_, *n*_*d*_, *t*_*Fc*_, *T*_*T*_) and presentation of a general model to calculate total frame time *T* with an *a priori* unknown variable overhead time *t*_*o*_. From a measurement of *T, t*_*o*_ can be calculated with *t*_*o*_ = (*T* − *T*_*T*_)/(*n*_*F,x*_*n*_*F,y*_) − *t*_*Fc*_. (b) It turns out that *t*_*o*_ depends strongly on *n*_*F,x*_ and *n*_*F,y*_ for *n*_*F,x*_×*n*_*F,y*_ *<* 2000, but is largely constant for *n*_*T,x*_×*n*_*T,y*_ *>* 2000. Typically, *n*_*T,x*_×*n*_*T,y*_ ≥ 2500, so basically no dependence is assumed. This curious increase of the overhead for a low number of force map pixels is most likely caused by the digital oscilloscope used for data collection operated in so-called “rapid block mode”, which is designed to reduce the delay when collecting many data blocks (e.g. force curves). However, when the number of data blocks is low, the overhead might become noticeable. (c) There is a slight dependence of *t*_*o*_ on *n*_*d*_, but only for *n*_*d*_ > 4000. Usually, *n*_*d*_ = 1000 due to limitations of disc space and, again, no dependence is assumed. (d) The topography pixel number *n*_*T,x*_ and *n*_*T,y*_ have almost no effect on *t*_*o*_. (e) Finally, the force curve time *t*_*Fc*_ and the topography time *T*_*T*_ both linearly increase the overhead *t*_*o*_. Using a multivariate linear fit, we can estimate *t*_*o*_ = 1.166 *ms* + 6.011 × 10^−2^*t*_*Fc*_ + 8.165 × 10^−5^*T*_*T*_, with *t*_*Fc*_ and *T*_*T*_ in milliseconds. Using this relation, it is possible to estimate the HS-iFM frame time *T* by *T* = *T*_*T*_ + (*t*_*Fc*_ + *t*_*o*_)*n*_*F,x*_*n*_*F,y*_. It should be noted that this estimation will vary with the specific system configuration, and is expected to be especially influenced by the data collection equipment. Here, a PicoScope 5443D (Pico Technology, UK) was used. However, the analysis procedure presented in this figure can be applied to the characterization of any system.

**Figure S3:**
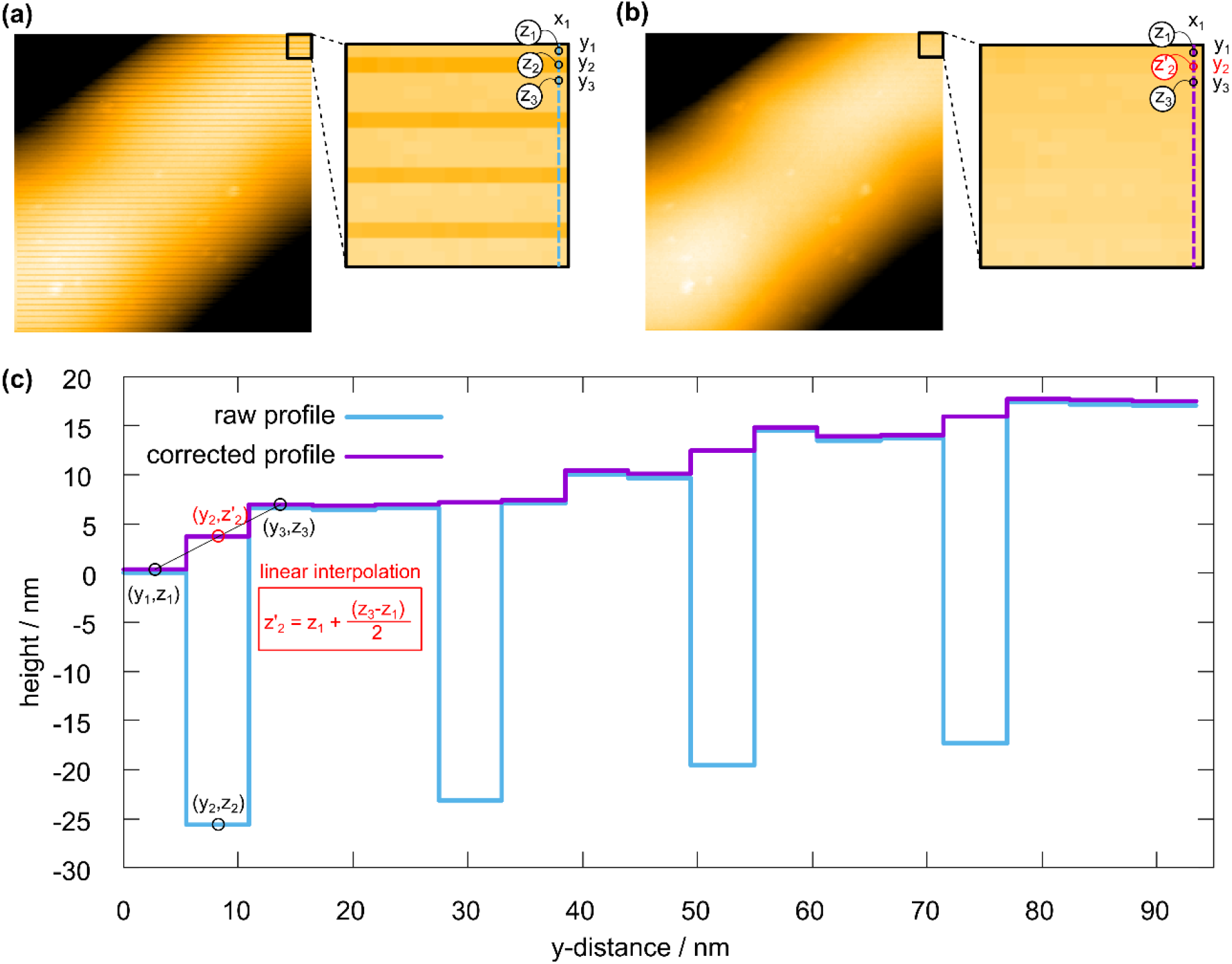
Removal of stripes caused by force mapping. (a) The dark horizontal lines are caused by the force mapping procedure, most likely because the PID-controller is overwhelmed by rapid switching between topography and force curves in these lines. The zoom clearly shows the regular-spaced lines. (b) The same image after line removal. The zoom shows that the horizontal lines were successfully removed (c)Line profiles in y-direction with constant *x*-coordinate *x*_*1*_ along the blue and purple dashed lines in (a) and (b), respectively. The line removal is done by linear interpolation between two adjacent pixels, as is demonstrated in (c).

**Figure S4:**
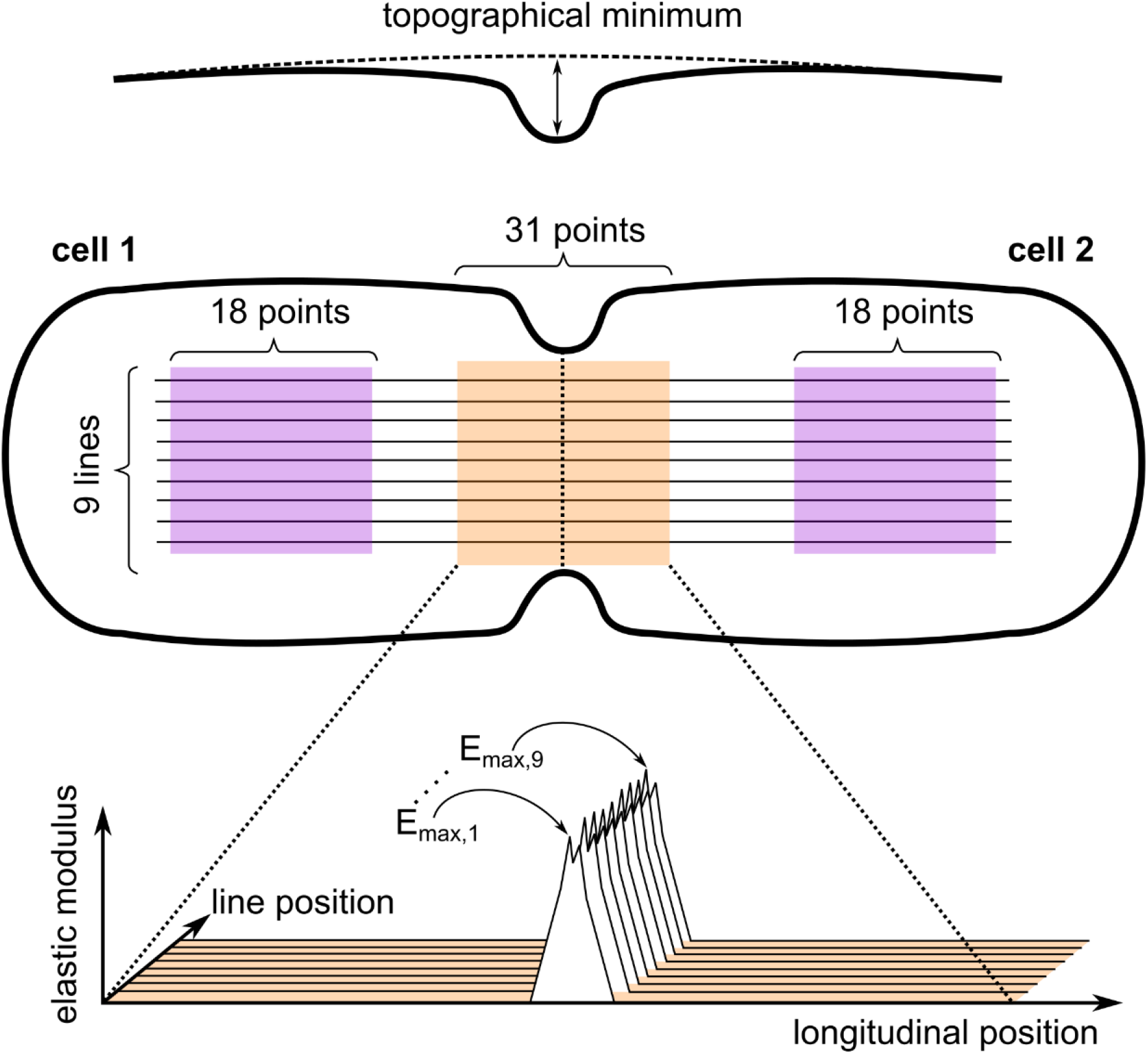
Scheme of data selection for contact mechanical analysis. 9 lines were drawn across the cell so that the division site was roughly centered. Along these 9 lines topography profiles and elastic modulus profiles were extracted. First, the topographical minimum is determined. Then, the maximum of the elastic modulus is found within 31 points centered around the topographical minimum. The average modulus of the division site is then calculated as the mean value of the nine maximum values. Finally, the average modulus of the left and right cells (cell 1 and cell 2) is determined as the mean of 162 modulus values (18 points × 9 lines).

**Figure S5:**
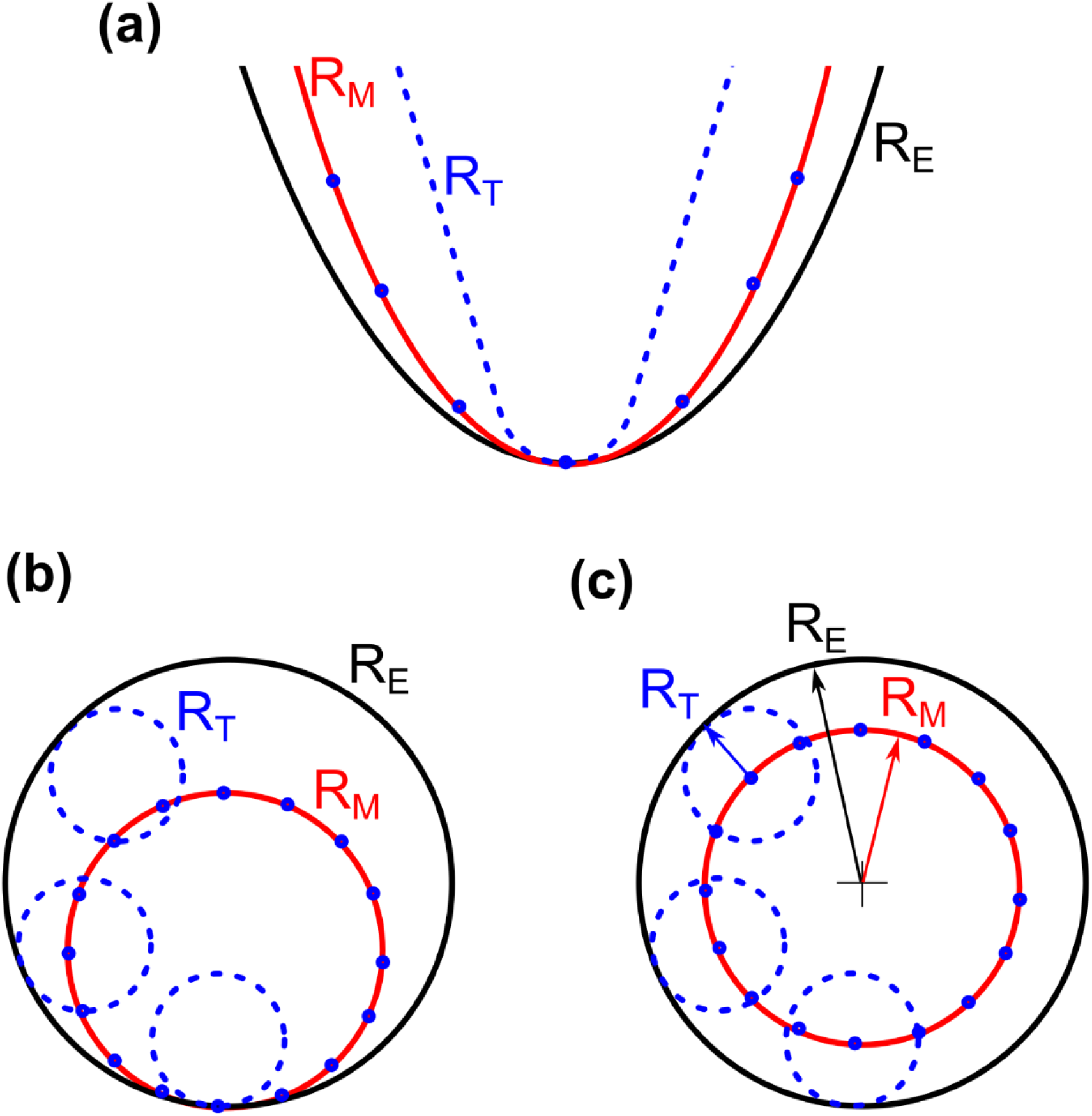
Tip-sample dilation effect. (a) A tip with radius R_T_ (blue dashed lines) is tracing a curved surface (black solid lines) with apex radius R_E_, resulting in the measured trajectory (red solid lines) with measured (apparent) radius R_M_. The tip apex is used to trace the measured trajectory (blue circles). (b) A simplified version of (a) using a small circle with radius R_T_ to trace a larger circle with radius R_E_ resulting in a measures circle with radius R_M_ = R_E_ – R_T_. (c) Shifting the circle’s tracing point to the center, it becomes clear that R_M_ = R_E_ – R_T_. Thus, the true concave radius R_E_ of a sample can be estimated by *R*_E_ = *R*_M_ + *R*_*T*_.

**Figure S6:**
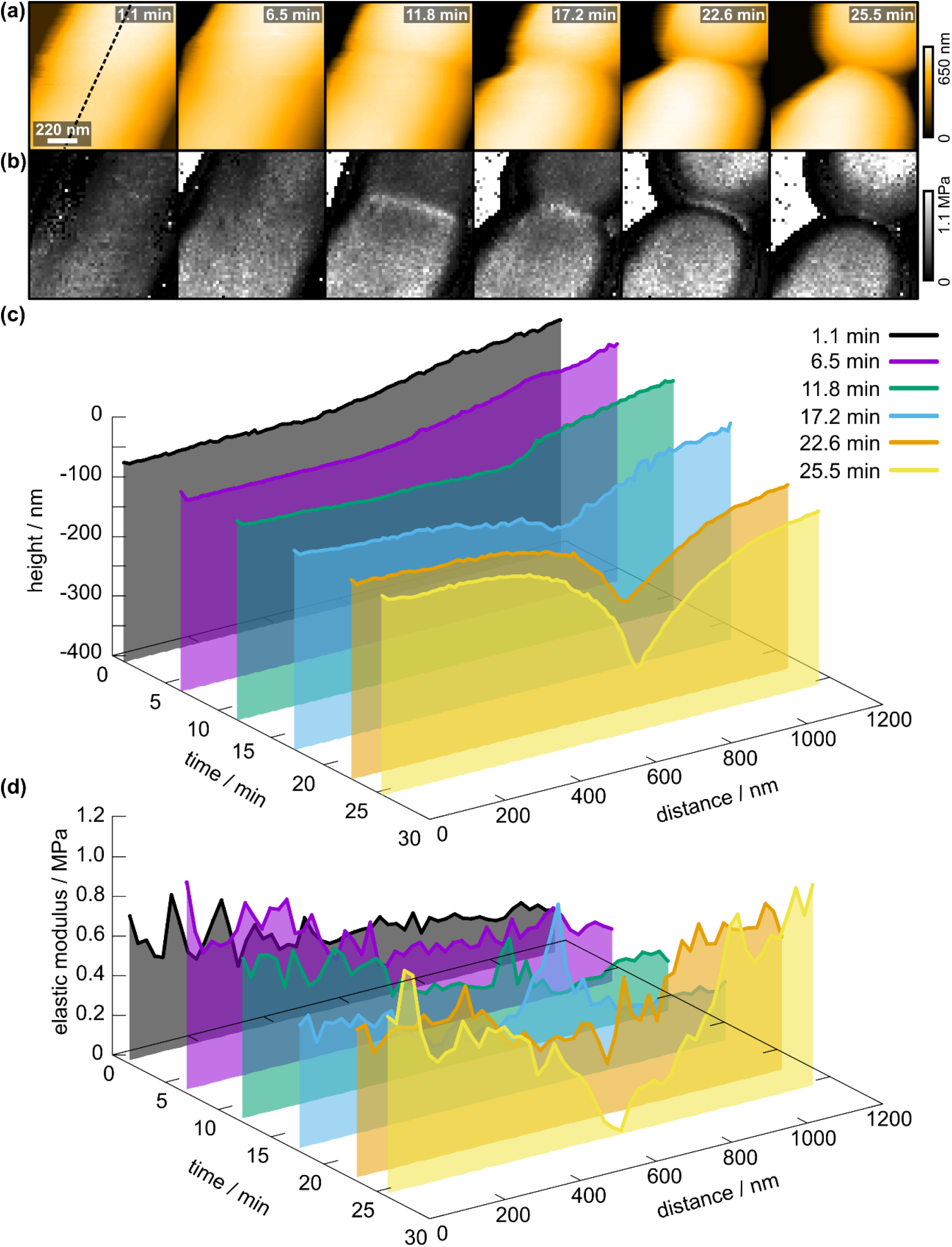
Details of topography and elastic modulus changes during cell division. Maps of (a) topography and (b) elastic modulus recorded on the same cell as main text Figure 2. Line profiles of (c) topography and (d)elastic modulus highlight the considerable changes around the division site. The position of the line profiles is marked with a dashed black line in (a).

**Figure S7:**
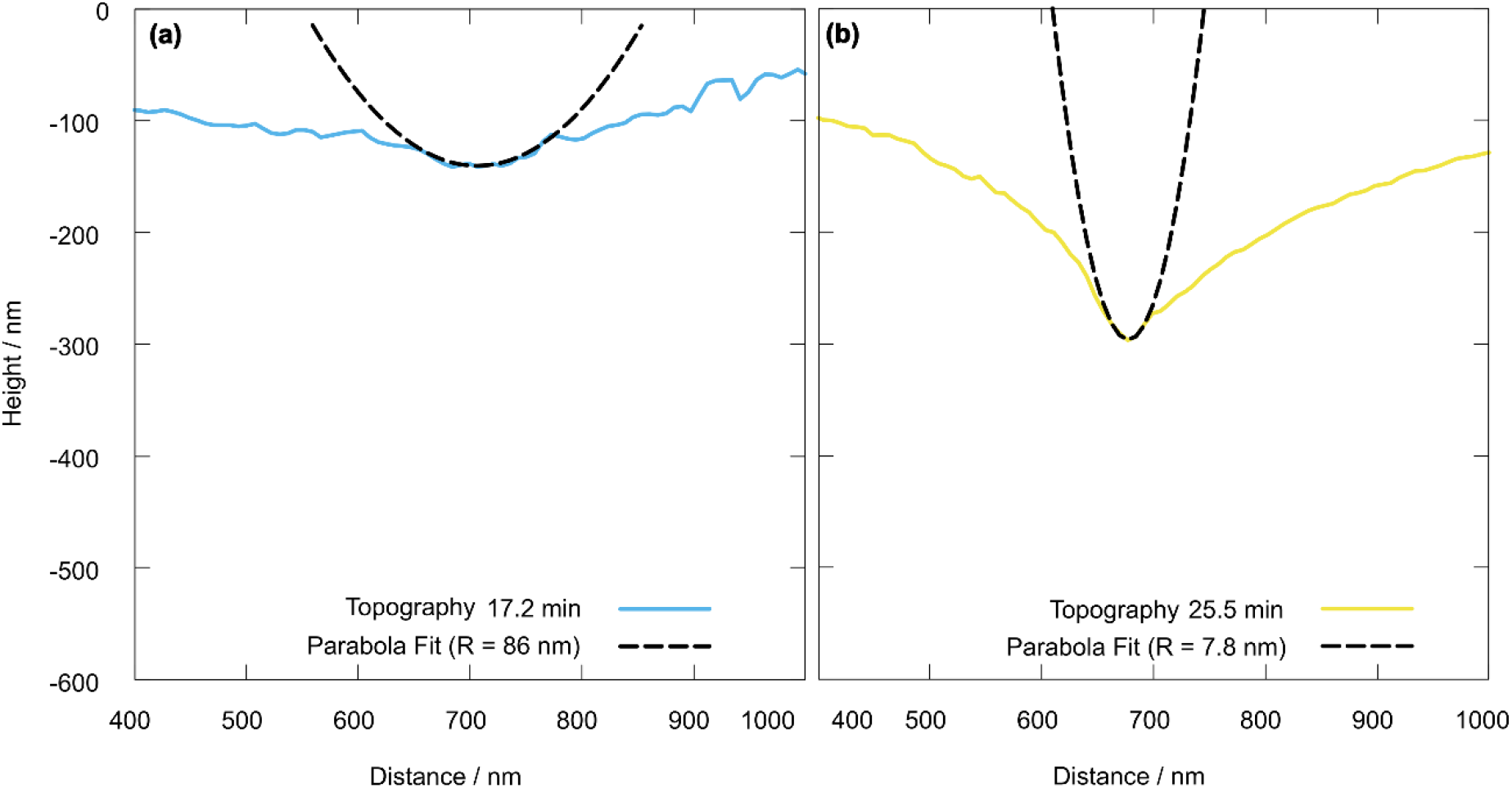
Representative fitting of the longitudinal radius at the division site with a parabola at (a) 17.2 min and (b) at 25.5 min. The corresponding topography and elastic modulus maps are shown in Supplementary Figure S6. This fitting procedure is done for 9 lines across a single frame and repeated for all frames. The longitudinal radius designated for a single frame is the minimum radius measured across the 9 lines. Note that before the division site becomes measurable the fitted radius typically corresponds to a small dent in the surface and has nothing to do with division. It is, however, a good indicator to see if other surface features will influence the elastic modulus measurement significantly. The results suggest that there is little influence.

**Figure S8:**
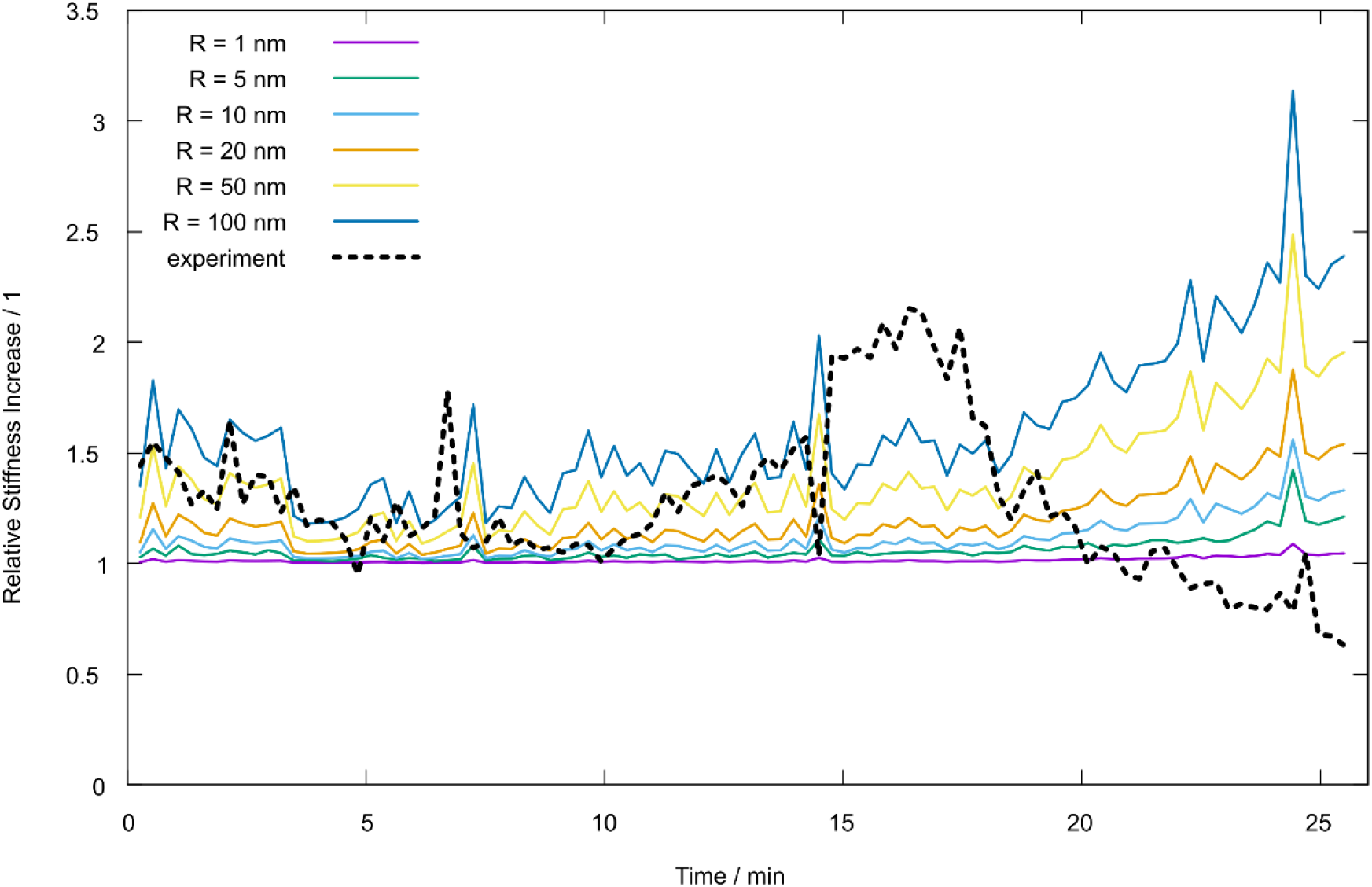
Influence of assumed tip radius on the apparent increase of the elastic modulus using the data from Figure 4b (main text). Even a tip radius of 100 nm cannot explain the rapid increase of stiffness in the division area. It should be noted that it is unlikely that a tip with a radius of 100 nm used. Typical tip radius values range from 1 nm to 10 nm.

**Figure S9:**
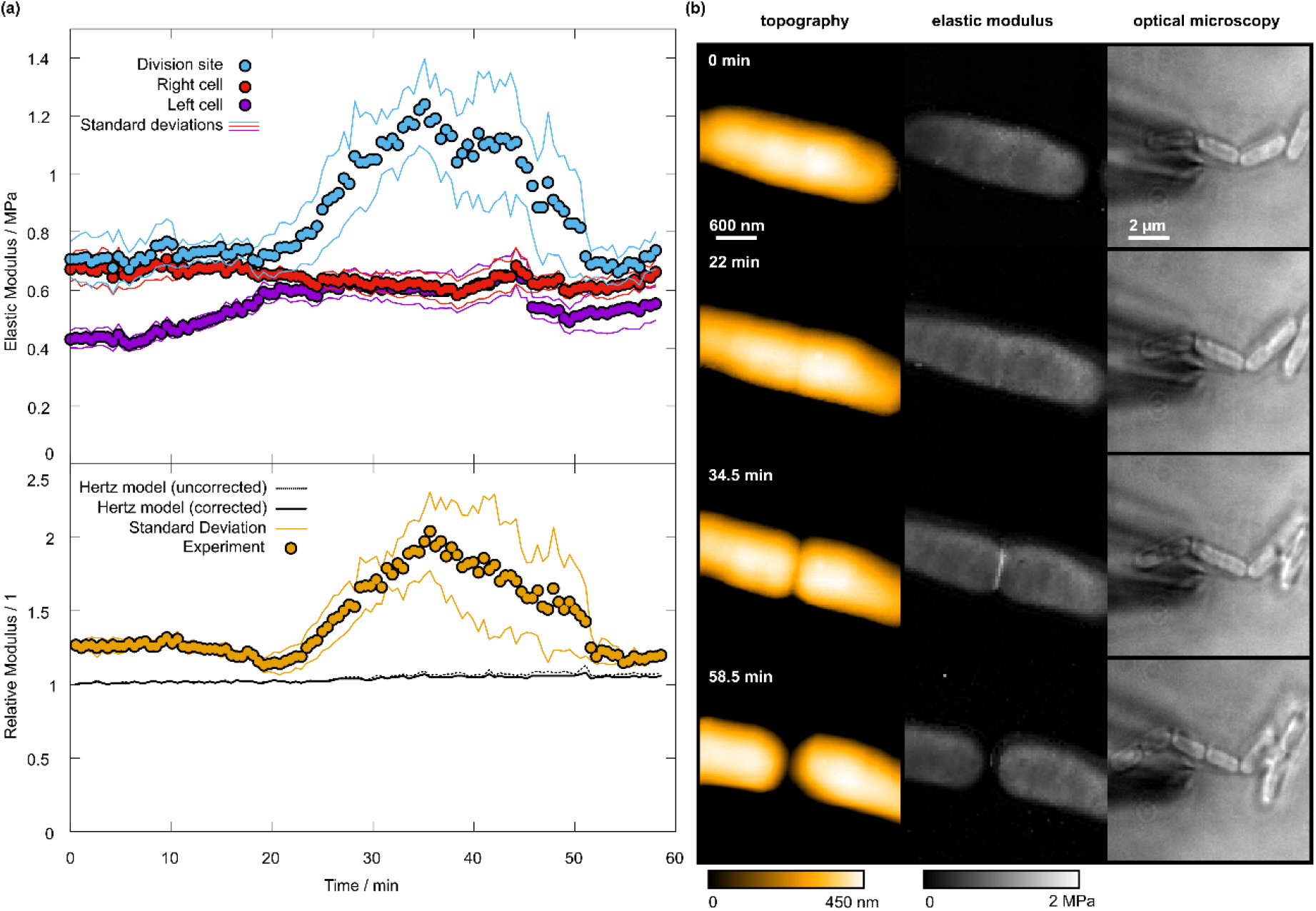
Mechanical properties and topography of *E. coli* division on a 3 µm × 3 µm scale. (a) Temporal evolution of the elastic modulus of the left and right cells as well as the division site (top graph). The bottom graph depicts the relative increase in elastic modulus of the division site with respect to the undeformed areas. The black lines show the theoretically expected elastic modulus increase of the division site by topography. The solid line is calculated from the tip-dilation corrected topography, while the dashed line is calculated from the uncorrected topography. (b) Snapshots of the division process as topography maps (left), elastic modulus maps (center), and optical microscopy (right).

**Figure S10:**
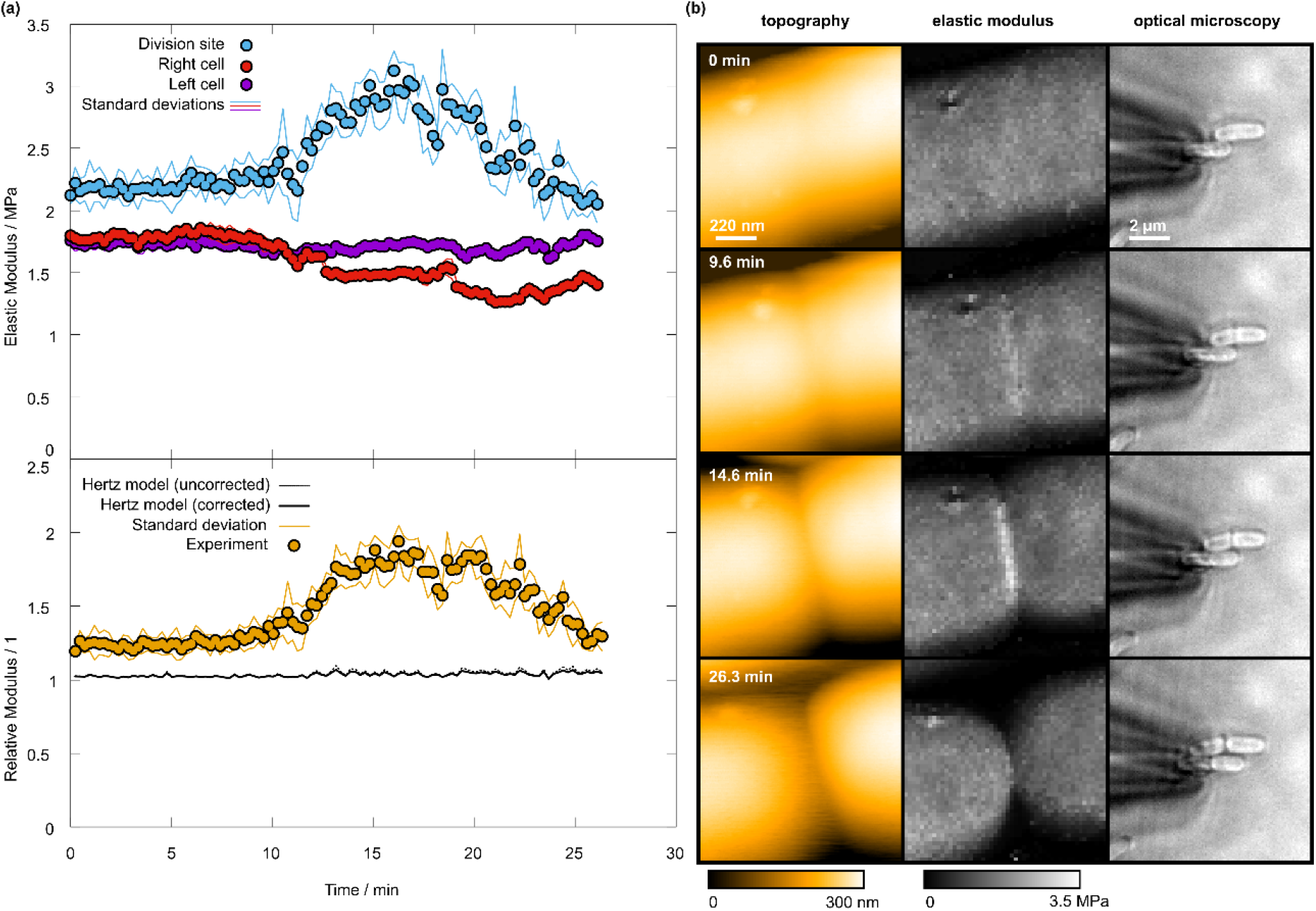
Mechanical properties and topography of *E. coli* division on a 1.1 µm × 1.1 µm scale. (a) Temporal evolution of the elastic modulus of the left and right cells as well as the division site (top graph). The bottom graph depicts the relative increase in elastic modulus of the division site with respect to the undeformed areas. The black lines show the theoretically expected elastic modulus increase of the division site by topography. The solid line is calculated from the tip-dilation corrected topography, while the dashed line is calculated from the uncorrected topography. (b) Snapshots of the division process as topography maps (left), elastic modulus maps (center), and optical microscopy (right).

**Figure S11:**
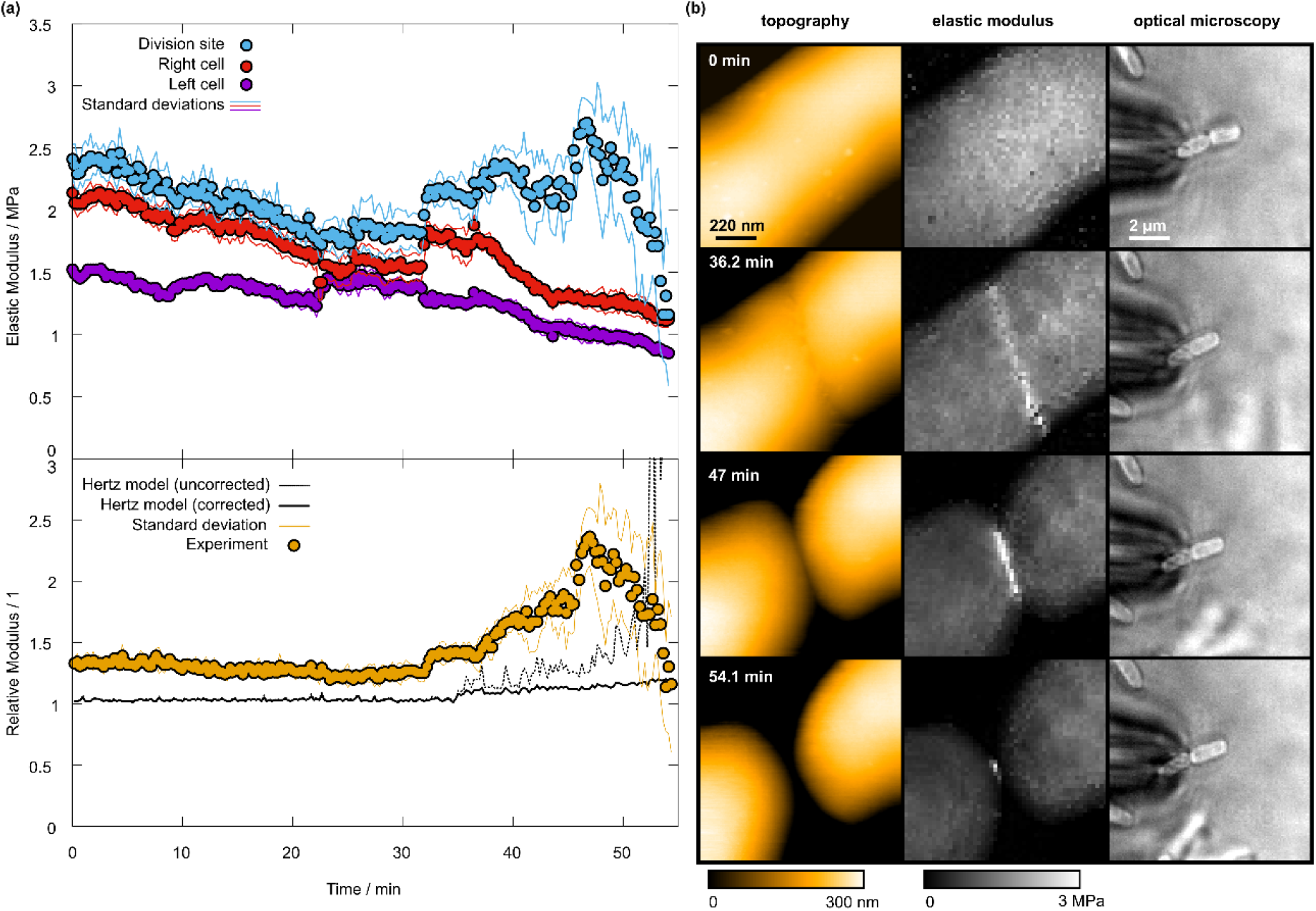
Mechanical properties and topography of *E. coli* division on a 1.1 µm × 1.1 µm scale. (a) Temporal evolution of the elastic modulus of the left and right cells as well as the division site (top graph). The bottom graph depicts the relative increase in elastic modulus of the division site with respect to the undeformed areas. The black lines show the theoretically expected elastic modulus increase of the division site by topography. The solid line is calculated from the tip-dilation corrected topography, while the dashed line is calculated from the uncorrected topography. (b) Snapshots of the division process as topography maps (left), elastic modulus maps (center), and optical microscopy (right).

**Figure S12:**
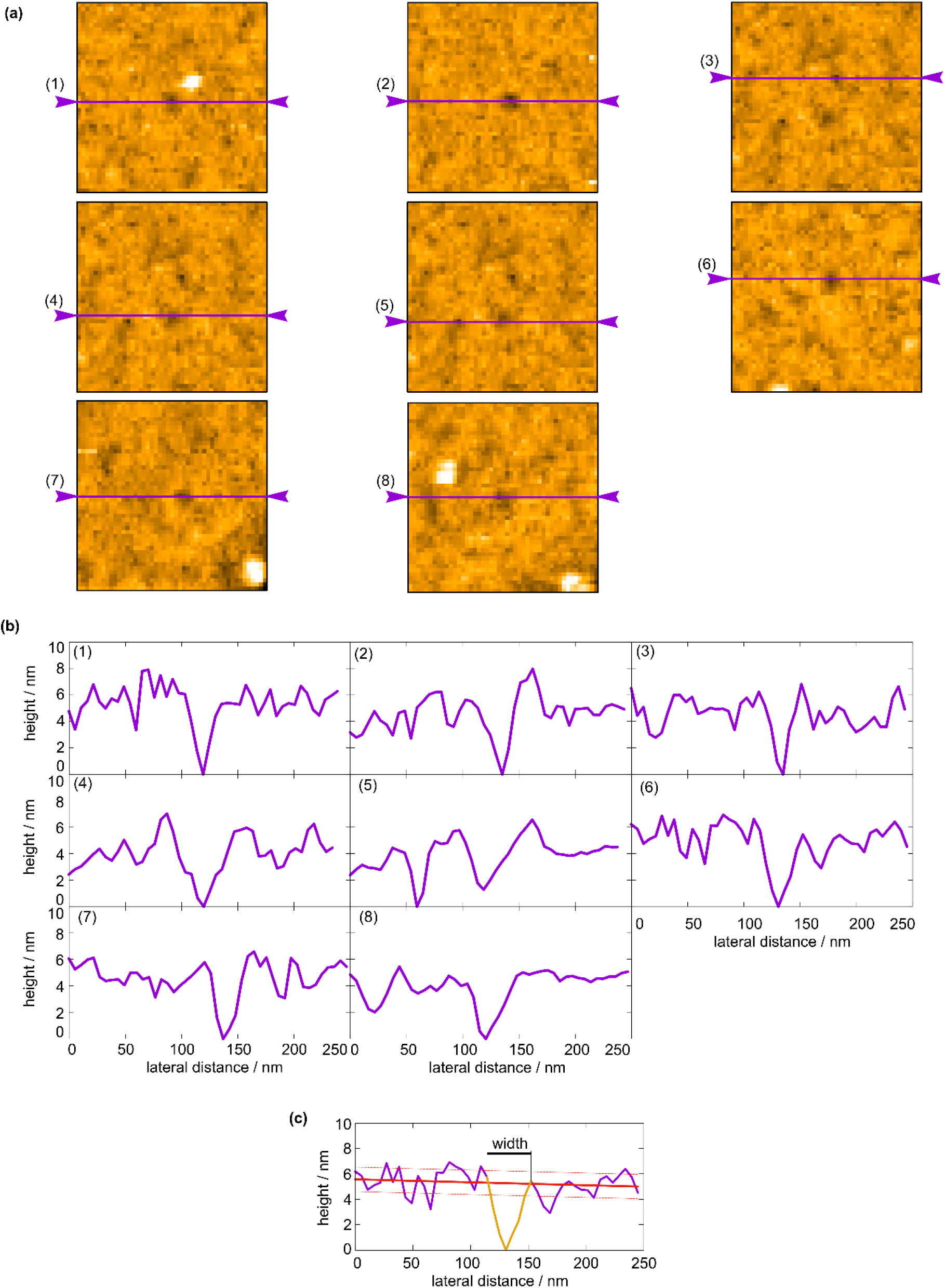
Pore size determination. (a) HS-AFM topography images of the 8 manually selected pores. (b) Manually placed line profiles of the pores as indicated by the purple arrows and lines in (a). (c) Exemplary of the automated pore size determination. First the minimum of the pore is determined and then by successively moving stepwise left until the lower range of the background (lower dashed red line) is reached. This is the left point of the hole. The same is repeated for the right side. The difference between these points is then the width (solid black line). The background is determined by a linear fit.

**Figure S13:**
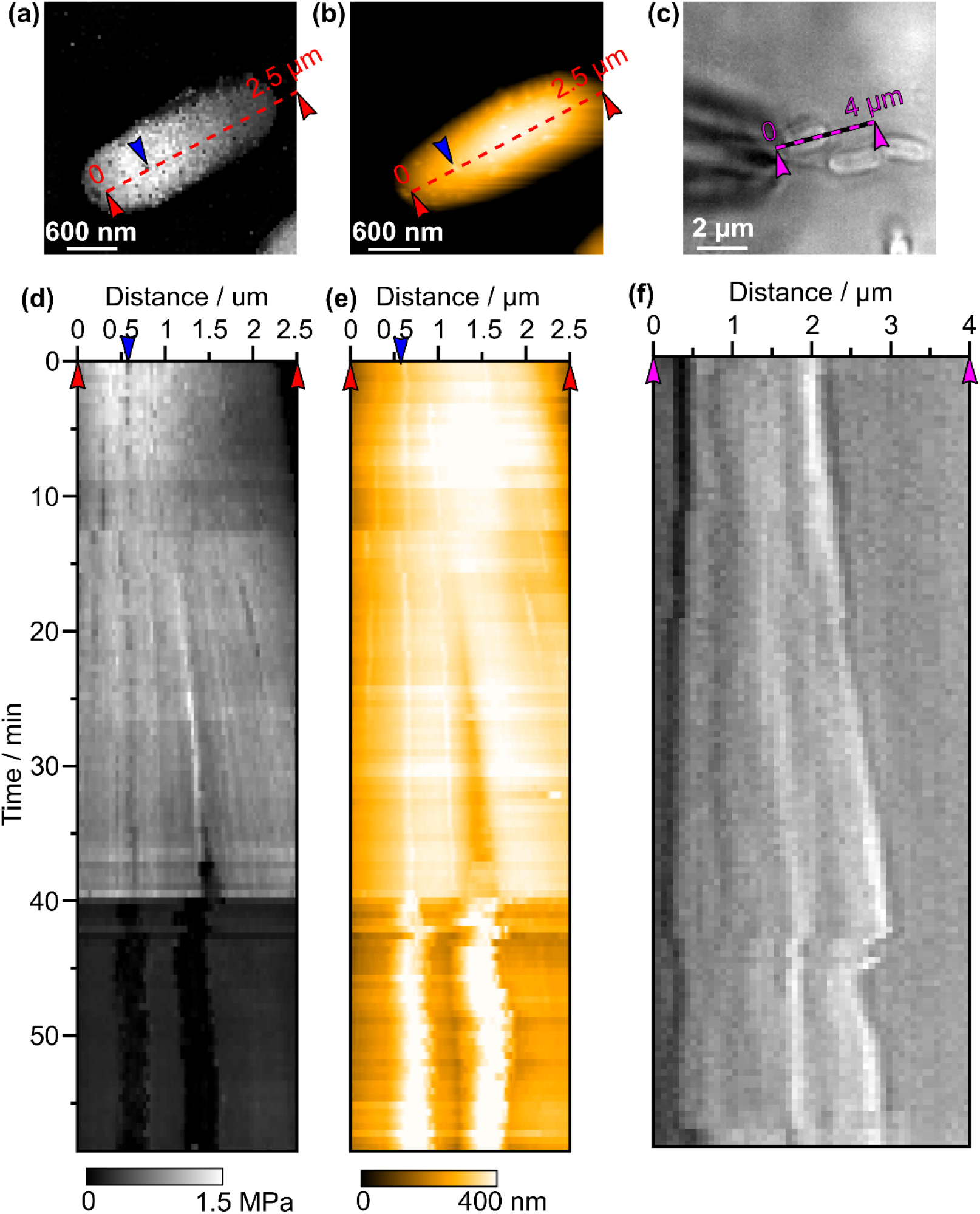
Kymograph of bursting cell in Figure 6 (main text). Position of the line profile in (a) Elastic modulus map, (b) topography, and (c) optical microscopy image. The profile in (a) and (b) were placed on the same positions. Kymographs of (d) elastic modulus and (e) topography along the line profile in (a), respectively (b). (f) Kymograph along the profile in (c). The red arrows in in (a), (b), (d), (e) demark the beginning and end of the line profile while the blue arrows mark the bursting vesicle. The purple arrows in (c)and (f) mark the beginning and end of the line profile. Due to drift and the cell growing, the position of the line profile in (a), (b), (d), (e) was manually adjusted to always include the bursting vesicle. At the 40 min mark the modulus drastically drops, which is concurrent with the bursting of the vesicle and a reduction in height. The latter could be caused by the deflating cell, which can be seen in (f) where the cells instantly shorten after the bursting event.

**Figure S14:**
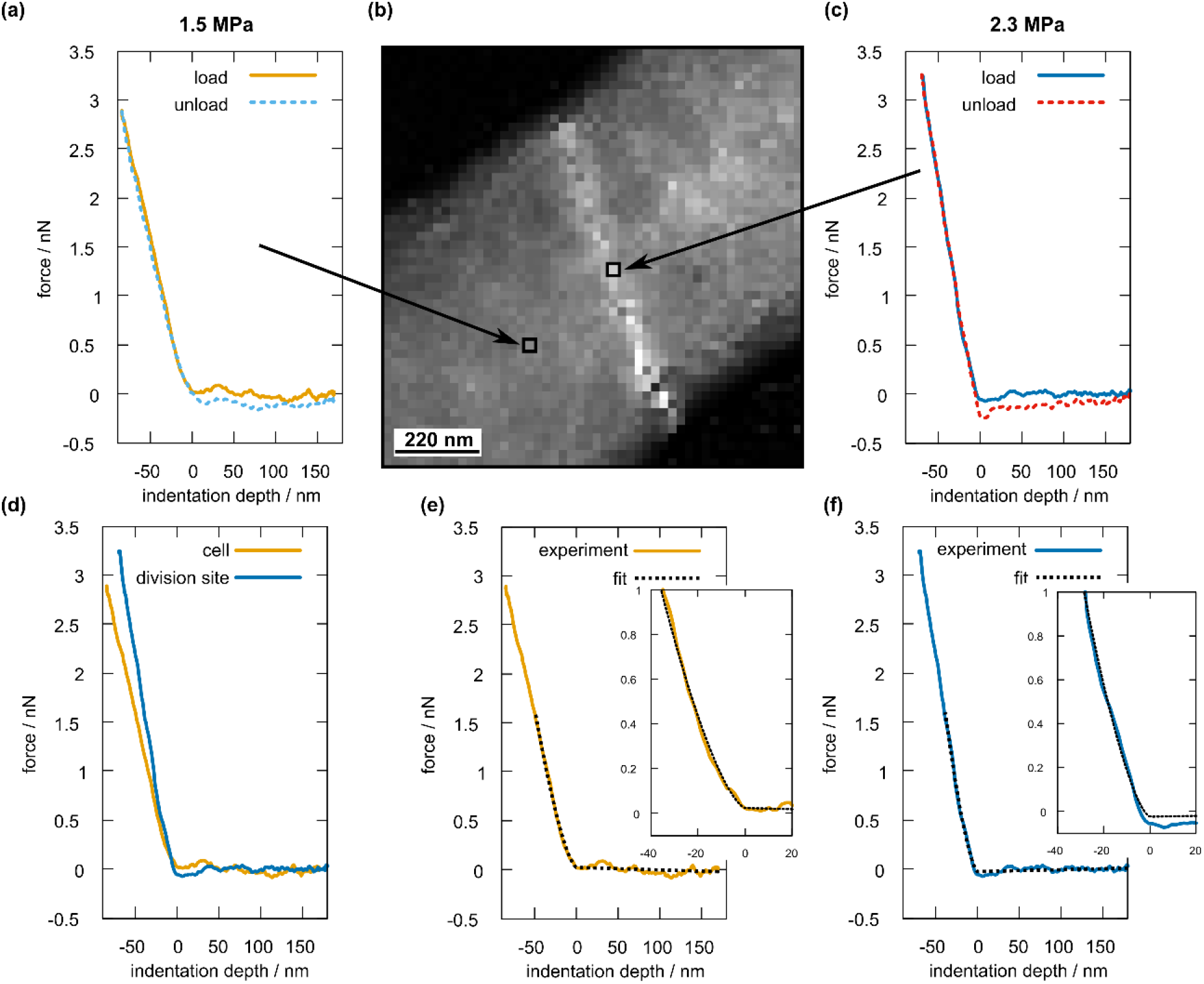
Exemplary force curves on a dividing *E. coli* cell. (a) Force curve recorded on the undeformed cell surface. (b) Force map (scale 0 – 3 MPa). (c) Force curve on the division site. (d) Comparison between the loading segments of (a) and (c). Hertz contact mechanics fits of the loading segments of (e) the undeformed surface and (f) the division site. The insets show zoomed-out sections of the fitted curves to demonstrate the quality of the fits.

**Figure S15:**
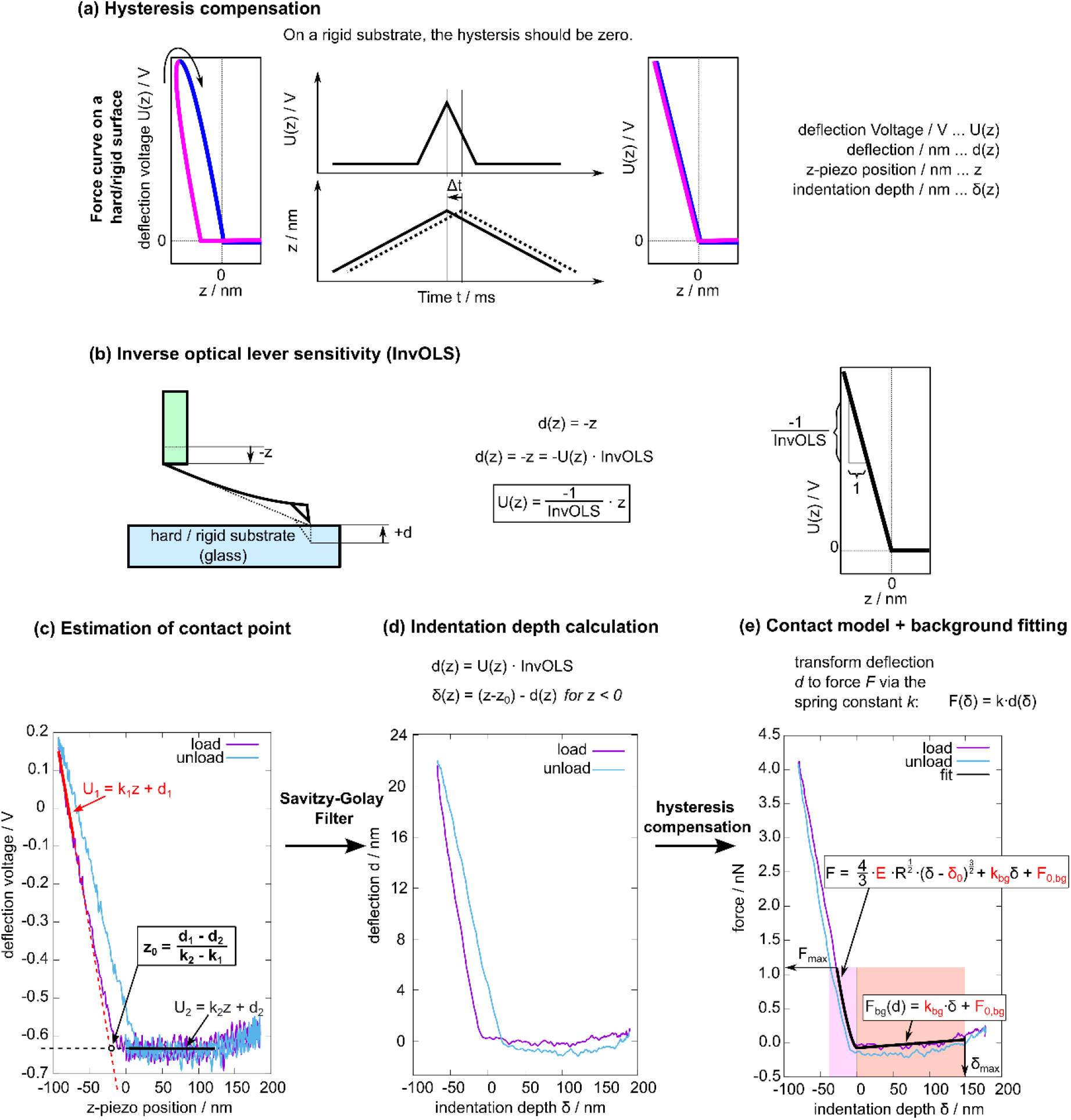
Scheme of the analysis procedure. (a) A lag between the piezo and the applied voltage will cause a hysteresis even when performing force curves on rigid substrates. This is used to calibrate the lag. Shifting the *z*-piezo distance to earlier times until the hysteresis is zero allows to determine the lag *Δt*, which is then used to shift all curves. (b) The inverse optical lever sensitivity (InvOLS) is also determined on a hard substrate after the hysteresis was compensated for. (c) Initially, the contact point *z*_*0*_ is determined by fitting the load curve from both ends and setting the crossing as *z*_*0*_. (d) After applying a Savitzky-Golay filter to remove the noise, the *z*-piezo distance is converted to indentation depth using the deflection. (e) After hysteresis compensation and converting deflection into force, the curve is fit with a Hertz contact model with variable zero-point and an additive linear background. The variables in red are the fit parameters. The fitting region is limited by the maximum force *F*_*max*_ and the maximum distance *d*_*max*_, both of which are manually chosen.

## Notes

### 1. Details of high-speed in-line force mapping

Relevant signals during HS-iFM are presented in Supplementary Figure S1a. The deflection signal shows the cantilever oscillating at approximately 500 kHz at the beginning (I in Supplementary Figure S1a and b), followed by the force application and continued with oscillation. The amplitude clearly depicts the phases of topography measurement (amplitude in the high state) and force curve recording (amplitude in the low state, II in Supplementary Figure S1a and b). The PID hold signal is responsible for switching the HS-AFM’s proportional-integral-differential (PID) controller from active to the “hold” state where the cantilever’s position is held constant and not changed by the controller. The *z*-piezo signal shows the waveform that is used to actuate the cantilever along the *z*-position. When the *z*-piezo signal reaches its minimum value, the deflection has its maximum (III in Supplementary Figure S1a and b).

The relationship between deflection and z-piezo signal is detailed in Supplementary Figure S1c-e. When force curve acquisition starts, the tip is close to the surface (i) and first is retracted as far as possible (ii). Then, the z-piezo extends towards the initial position (iii) and surpasses it, leading to an increase in the deflection signal due to the tip contacting the surface (iv). After the maximum z-piezo extension (v) is reached, the movement is reversed until the initial state is reached.

For each position on the force map grid, the recording of deflection and z-piezo voltages are triggered to happen simultaneously with the output of the z-piezo voltage. Both recording and output of z-piezo voltage are performed by an external, programmable digital oscilloscope (PicoScope 5443D, Pico Technology, UK). The oscilloscope features a bandwidth of 100 MHz and an internal memory of 128 MS, operated at a signal resolution of 15 bit. The z-piezo waveform can be freely adjusted in shape and length, but for the purpose of this study it was set-up as follows: 0.5 ms to retract from the surface to 2 V, corresponding to about 185 nm, in a sinusoidal shape, moving the cantilever as far away from the surface as possible. Next the cantilever is extended within 0.5 ms to a maximum extension value that is chosen individually for each sample and the desired forces. This is followed by a symmetric retraction and the repositioning to the initial position (0 V piezo extension) in 0.5 ms. The overall time is thus 2 ms per force curve. However, as can be seen in Supplementary Figure S1a, the time of the PID controller set to “hold” is about 3 ms, which is caused by overheads of settings for the scanning procedure and the oscilloscope, which need to be completed before switching the PID controller on and scanning can continue. In order to estimate the time an HS-iFM frame will take before the start of a measurement, a closer investigation is necessary to identify the dependence of frame time on each scanning parameter. The detailed results are highly system specific and are summarized in Supplementary Figure S2. Using the above force curve time schedule, a 50×50 HS-iFM force map will require 15 s to record, while a 75×75 force map requires about 30 s, including 10 s for topography recording.

Note that there is also no hard limit on the maximum force map pixel resolution, but since each force map pixel will contain one force curve, disk space may be a limiting factor. For example, a 50 × 50 force map with individual force curves containing 1000 datapoints (each point is represented as a 16-bit integer value), 100 HS-iFM frames will occupy approximately 1 GB of disk space.

Finally, when using HS-iFM, stripes are introduced in the topography images. The reason for this is the rapid switching of the PID controller as well as the cantilever *z*-position movement for hundreds of nanometers during a force curve. This results in horizontal stripes in the topography scan – or, more precisely, in a step between a pure topography line and a force curve containing line. To further process the images, these stripes need to be removed, which is done by interpolating the distorted line from the preceding and succeeding lines (Supplementary Figure S3). Such interpolation is a simple and effective method for de-striping that can be employed due to knowing the exact positions of the stripes beforehand, making the use of more complex de-striping algorithms unnecessary.

### 2. Contact mechanics with elliptical contact area

The general theory can be simplified to express the force as a function of indentation depth *δ* and eccentricity *e*:

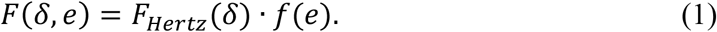

With *F*_*Hertz*_ describing the Hertzian contact mechanics for circular contact areas in the well-known form as

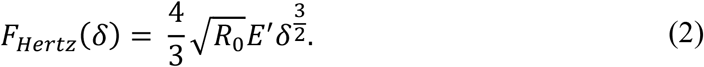

R_0_ is the effective radius of two spheres in contact, generally, and defined as *R*_0_ =(1/*R*_*t*_+ 1/*R* _*s*_)^−1^, where *R*_*t*_ is the tip radius and *R*_*s*_ is the sample’s radius. *E’* is the reduced elastic modulus given as 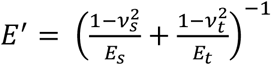. *E*_*s*_ and *E*_*t*_ denote the elastic modulus of the sample and tip, respectively, while *v*_s_and *v*_t_are the respective Poisson’s parameters. For our purpose we will assume to be *R*_*s*_ *>> R*_*t*_and thus *R*_*0*_will simplify to *R*_*0*_ *≈ R*_*t*_. Furthermore, since we can expect that the tip (electron deposited carbon) is much stiffer than the sample 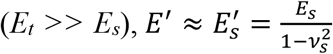. Thus, the general equation (2) changes to the specific relation

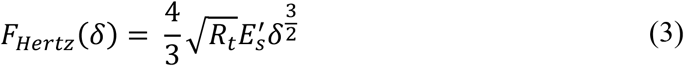

Equation (3) describes the contact mechanics of a rigid spherical tip indenting a flat, infinitely extended, elastic plane, as is commonly the case for many applications.

The remaining factor in equation (1), *f(e)*, is the correction factor for the elliptical contact area and is dependent on the contact area’s eccentricity *e*. The eccentricity *e* has to be determined numerically from the relation

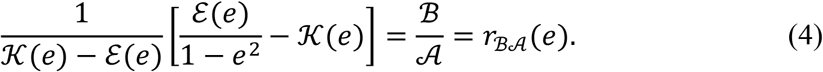

The functions 𝒦(*e*) and *ℰ*(*e*) are the first and second complete elliptical integrals, respectively. 𝒜 and ℬ are the principal curvatures in *x* and *y* directions, respectively and are calculated from the radii of tip *R*_*tx*_ *= R*_*ty*_ *= R*_*t*_, and sample *R*_*sx*_ and *R*_*sy*_ as

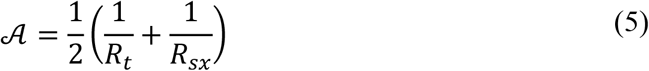

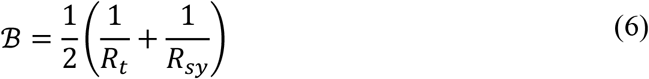

Note that the eccentricity *e* depends only on the ratio ℬ/𝒜. Thus, numerical estimation of *e* can be achieved by calculating *r*_ℬ𝒜_(*e*) successively from an arbitrary starting point until *r*_ℬ𝒜_(*e*) ≈ ℬ/𝒜. Another possibility is to plot *r*_ℬ𝒜_(*e*) and select the value of *e* corresponding to *r*_ℬ𝒜_(*e*) ≈ ℬ/𝒜 [53]. Since equation (4) will only hold for ℬ/𝒜 > 1, the coordinate system should always be chosen so that ℬ > 𝒜, meaning that the *y* axis should be aligned so that it coincides with the elliptical contact area’s minor axis and the *x* axis with the major axis.

Finally, the correction factor for the elliptical contact area can be determined as

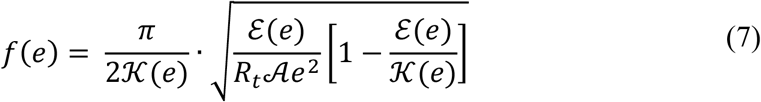

For circular contact areas *e* = 0 and 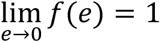, which can be verified numerically by plotting *f(e)*. This means that *F*(*δ, e*) naturally collapses into the classical Hertzian theory for circular contact areas.

